# LinRace: single cell lineage reconstruction using paired lineage barcode and gene expression data

**DOI:** 10.1101/2023.04.12.536601

**Authors:** Xinhai Pan, Hechen Li, Pranav Putta, Xiuwei Zhang

## Abstract

Understanding how single cells divide and differentiate into different cell types in developed organs is one of the major tasks of developmental and stem cell biology. Recently, lineage tracing technology using CRISPR/Cas9 genome editing has enabled simultaneous readouts of gene expressions and lineage barcodes in single cells, which allows for the reconstruction of the cell division tree, and even the detection of cell types and differentiation trajectories at the whole organism level. While most state-of-the-art methods for lineage reconstruction utilize only the lineage barcode data, methods that incorporate gene expression data are emerging, aiming to improve the accuracy of lineage reconstruction. However, effectively incorporating the gene expression data requires a reasonable model on how gene expression data changes along generations of divisions. Here, we present LinRace (**Lin**eage **R**econstruction with asymmetric cell division model), a method that integrates the lineage barcode and gene expression data using the asymmetric cell division model and infers cell lineage under a framework combining Neighbor Joining and maximum-likelihood heuristics. On both simulated and real data, LinRace outputs more accurate cell division trees than existing methods. Moreover, Lin Race can output the cell states (cell types) of ancestral cells, which is rarely performed with existing lineage reconstruction methods. The information on ancestral cells can be used to analyze how a progenitor cell generates a large population of cells with various functionalities. LinRace is available at: https://github.com/ZhangLabGT/LinRace.

## 1 Introduction

Understanding how cells divide and differentiate into various cell types is a fundamental problem in developmental biology. *Lineage tracing* technology which traces cell divisions using a “recorder” is the most widely used technique to study the developmental histories of cells, while traditional lineage tracing technologies can only work with a limited number of cells with low resolution [16]. Recently, sequencing-based lineage tracing methods (e.g. using CRISPR/Cas9 genome editing) have enabled the simultaneous recording of the clonal relationships of single cells alongside the transcriptomes [36] for up to thousands of cells. Such methods utilize lineage recorders, which are exogenous DNA sequences integrated into the genome. Even though there are different ways of designing the lineage recorder [1,2,4,13,20,25,31,33], the common idea is to introduce changes at the target sites (the location to which the Cas9 protein binds to induce mutations) on the lineage recorder which accumulates through generations of cell divisions. Finally, the recorders are sequenced together with the transcriptome of every single cell, resulting in the “lineage barcode” data. Each target site corresponds to a character in the barcode. The barcode data are used to reconstruct the cell division tree, which is also called the cell lineage tree in this paper. The reconstructed cell division tree can shed light on the developmental process that can not be directly measured.

However, inferring the cell division tree of a massive number of cells is a challenging problem. In addition to the computational complexity of inferring the lineage tree itself, the quality of the barcode data has posed a further challenge to this problem [30]. First, the number of target sites, which is the length of the string used for tree reconstruction, is usually small (the length of the string is 18 in a mouse embryo dataset [4] and 9 in a zebrafish dataset [25]); Second, dropouts in the CRISPR/Cas9 induced lineage barcode data can cause missing information in the data. There are two types of dropouts: one is called *excision dropout*, or collapse dropout [7], result in the loss of consecutive target sites in between two simultaneous mutations (in this case, the mutations are deletions in the barcode). The other type of dropout is due to the limited capture efficiency of the sequencing experiment, where the barcodes of certain cells are not profiled. Finally, the biased distribution of mutations across the barcode and the number of mutations, represented by the *mutation rate* parameter, may not be optimal for reconstructing the cell division tree. Is it shown that the distribution of mutations across the barcode is not uniform, but rather biased towards certain target sites [29,24]. The mutation rate, which represents the efficiency of mutations being induced in the barcode, has a major impact on the potential to successfully infer the cell division lineages from the data. However, current experimental technologies do not guarantee that the mutations occur at a rate that allows the tracing of every cell division event.

Various methods for tree inference from lineage barcodes have been developed. Recently, a DREAM challenge was held to gather the community effort to compare the state-of-the-art lineage tree inference methods [9]. Among the benchmarked algorithms the best performers are: DCLEAR [10], a distance-based method that first calculates the pairwise distance between cells and then reconstructs the cell lineage using bottom-up (agglomerative) algorithms such as Neighbor Joining (NJ) [28] or FastME [17]; Cassiopeia [12], a parsimony based method that aims at minimizing the number of mutations occurred on the reconstructed lineage tree. However, current methods for cell lineage tree reconstruction do not provide satisfying results using barcode data [9,24,29]. Moreover, due to the short barcode length and dropouts, a number of cells can have the same barcode, and the reconstructed lineage trees tend to have low depth (maximum path length from the root to a leaf node) and few internal nodes (with some nodes having large degrees) even using a perfect method. More recently, methods that combine lineage barcode and gene expression data are proposed, aiming to further improve the accuracy of cell lineage reconstruction. LinTIMaT [38] develops a combined likelihood function and uses a local search framework to search for the tree with the maximum likelihood. Integrating paired gene expression to infer the cell lineage tree can potentially refine the reconstruction, however, despite that the paired data should theoretically provide more information than barcodes alone, LinTIMaT did not beat methods that use only the lineage barcode data according to previous comparisons on synthetic datasets [24].

The key to combining the lineage barcode and gene expression data is to model the relationship between the two types of data, *i.e.*, how the gene expression of cells changes along with the barcode data during cell divisions. However, it is still an open question how the cell’s transcriptome changes during cell division. A simple assumption is to assume that cells that have similar transcriptomes should locate close in the cell division tree, *i.e.*, they should have similar barcodes. This is the assumption used in LinTIMaT. However, in a few recent papers that present paired single-cell lineage barcode and gene expression data, it is observed that, in a tree reconstructed using the barcode data, although a proportion of cells with the same cell state located in the same subtree, it is remarkable that some cells of the same cell state are located in different subtrees, and the same subtree can have multiple cell types [4,25]. We call this phenomenon the “partial consistency between transcriptome similarity and barcode similarity”.

The *asymmetric cell division model* has been shown to be able to account for the “partial consistency between transcriptome similarity and barcode similarity” [24]. It is commonly considered that cells can divide in a symmetric or asymmetric manner [15,21]. A symmetric cell division gives rise to daughter cells with the same cell state as the parent cells. During an asymmetric cell division, one daughter cell keeps the parent’s cell state, and the other one differentiates into a future cell state according to the cell state tree. For a given cell, there is a probability with which it divides asymmetrically, termed as the *asymmetric division rate*, denoted by *p_a_*. It has been shown that this probabilistic asymmetric cell division model leads to realistic paired single-cell lineage barcode and gene expression data.

To address these problems, we present LinRace, a new method that combines the lineage barcode and gene expression data to infer cell division trees, based on a joint Neighbor Joining (NJ) and maximum-likelihood framework. The asymmetric cell division model is used in LinRace to infer the states of ancestral cells and to calculate the likelihood function, thus incorporating the relationship between lineage barcode and gene expression data in a realistic way. On both simulated and real datasets, LinRace consistently outperforms the state-of-the-art methods according to multiple metrics. We show that the use of gene expression data in LinRace helps to improve the lineage tree reconstruction accuracy compared to methods that use only the lineage barcode data (Cassiopeia and DCLEAR). We also show that LinRace achieves better performances than the existing method that also uses gene expression data (LinTIMaT) while improving computational efficiency. Moreover, we demonstrate that when applied to large-scale real datasets, LinRace uncovers more detailed local lineage structures compared to LinTIMaT, as well as ancestral state information and state-lineage relationships that are consistent with observations from real data.

## 2 Results

### 2.1 Overview of LinRace

LinRace is motivated by the fact that currently available lineage barcode data cannot label each cell with a unique barcode, and can contain large sub-groups of cells with the same barcode. For example, the embryo2 dataset in the mouse lineage tracing system [4] has 19019 cells but only 2788 unique barcodes. Among all the barcodes, 1929 of the 2788 barcodes uniquely label one cell; 330 of the 2788 barcodes are shared by two cells. Supplementary Fig. 1a shows the number of cells sharing each barcode, and only the top 50 of the barcodes which are shared by the largest number of cells are shown. 86.4% of the cells share the same barcode with at least two other cells. The barcode data alone does not allow the inference of trees for the cells with the same barcode. LinRace aims to infer the cell lineage tree using paired lineage barcodes and gene expression data, where every single cell has its corresponding lineage barcode and gene expression data. The overview of LinRace is shown in Fig. 1a.

**Fig. 1.**
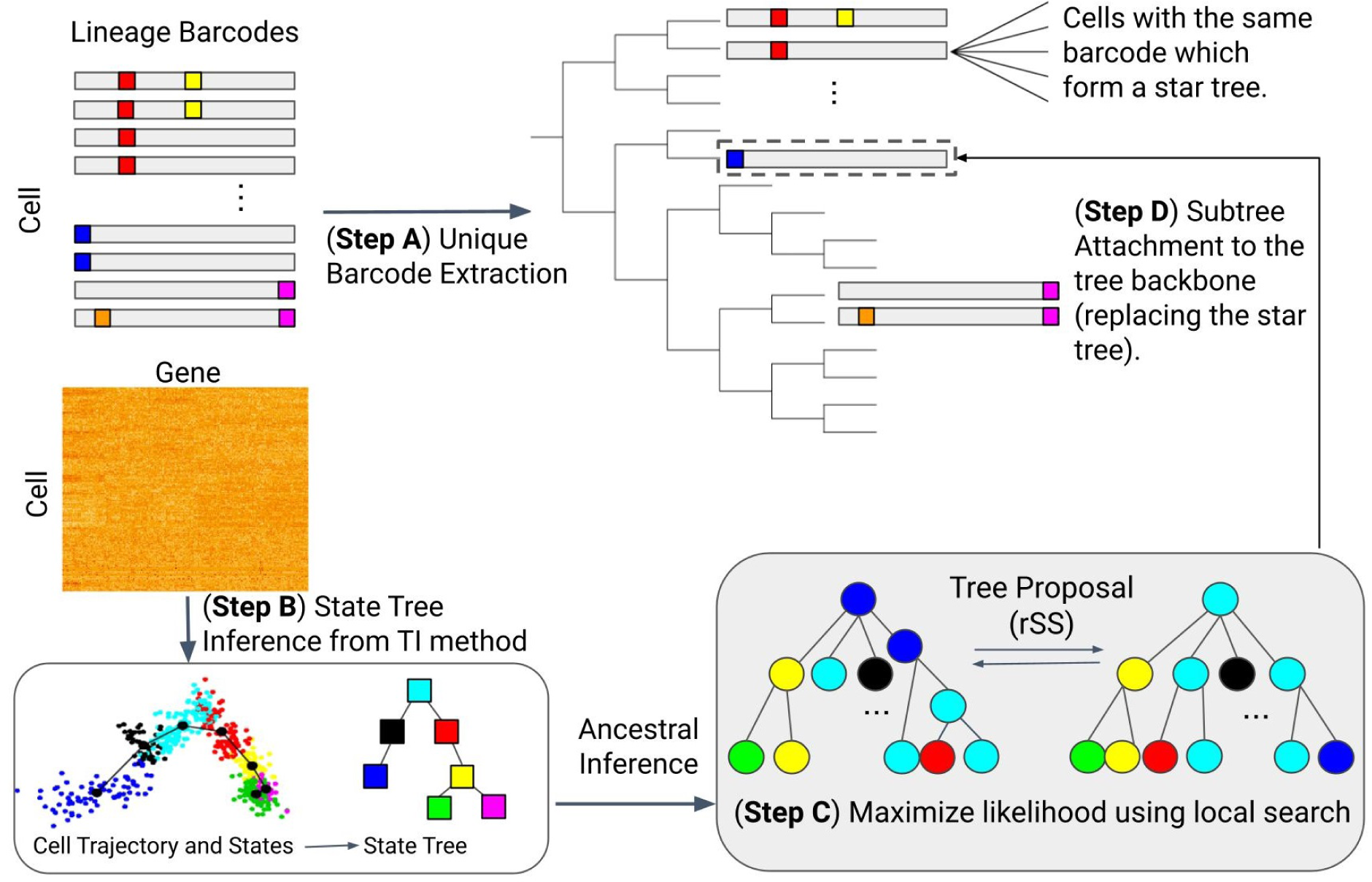
Overview of LinRace. **Step A** For barcode data, we extract the unique barcodes, and then perform Neighbor Joining to obtain the tree backbone, where each leaf represents a unique barcode shared by some cells. **Step B** For the gene expression data, we use K-means on PCA reduced dimensions to infer the cell states, and then use Slingshot[34] to infer the state trajectories which is then used to infer the ancestral states. **Step C** For each group of cells of the same barcode, we use a maximum likelihood + local search framework to find the subtree topologies of the cells. **Step D** The final output tree is obtained by combining the subtrees at their specific leaves on the tree backbone. Nodes in the trees that are illustrated by squares represent cell states (or cell types), and those illustrated using circles represent cells, and the color of each circle node represents the state of the cell.

First, from the barcode data of all the cells, we extract the unique barcodes and build a tree of these barcodes (Fig. 1 Step A) using the neighbor joining [28] tree reconstruction method. We then obtain a tree where each leaf node represents a unique barcode and can correspond to multiple cells with the same barcode. We denote this tree by *T*_0_. Then we use the single-cell gene expression data to refine the tree *T*_0_. We consider that there exists an underlying cell state transition mechanism that can be represented by a *cell state tree*, following which different cell types emerge during the cell division processes. This cell state tree can be inferred from the single cell gene expression data using trajectory inference methods [27,34,37] (Fig. 1 Step B). The cell state tree along with the *asymmetric cell division* model together model the relationship between lineage barcode and the gene expression data and allow for the inference of cell states for ancestral cells, as well as the design of the likelihood function used in Step C of Fig. 1. Details on the motivation for using the asymmetric division model are in Methods. Then, for each leaf in *T*_0_ which corresponds to one unique barcode and potentially many cells, we use a novel likelihood function (Methods) to find the best bifurcating tree (which means every internal node has exactly two children nodes) that maximizes this likelihood (Fig. 1 Step C). This likelihood is based on the asymmetric cell division model and the cell state tree. Finally, we attach the subtrees inferred in Step C to the lineage tree inferred from the barcode data (*T*_0_) to get the complete lineage of single cells (Fig. 1 Step D).

### 2.2 LinRace outperforms existing methods on synthetic datasets under various settings

We first test LinRace’s tree reconstruction performance against baseline methods using simulated data. We use TedSim [24] to generate simulated paired single-cell gene expression and lineage barcode data, which is the only existing simulator that generates such paired data with a ground truth cell division tree. We compare the results of LinRace with the state-of-the-art lineage tree reconstruction methods, including Cassiopeia-greedy and Cassiopeia-hybrid [12] (a parsimony-based method), and DCLEAR-kmer [10] (a distance-based method), which use only the barcode data, and LinTIMaT [38], a method based on the combined likelihood of gene expression and lineage barcode that use both types of data.

To obtain a comprehensive picture of the performances of the methods, we vary major parameters when generating the simulated data: (1) Mutation rate *µ*, the probability that a mutation (insertion or deletion) is induced per target site per cell division. Different mutation rates can result in barcode data with different quality, and it has been shown that the performance of lineage tree reconstruction methods using only the barcode data is affected by the mutation rate [24,29]. A range of [0.01, 0.3] was used in previous work [29], and we used a similar range which is [0.03, 0.3]. (2) With or without dropout. Real data contains a significant amount of dropouts, and simulated data with dropouts show properties (*e.g.* the distribution of the number of cells with the same barcode in Supplementary Fig. 1b) much closer to real data than simulated data without dropouts. We nevertheless test the methods on simulated data without dropouts which can show the effect of dropouts on each method. (3) Number of cells. The complexity of the lineage tree reconstruction problem increases with the number of cells. We generated datasets with 1024 and 4096 cells. For each combination of parameters, we generated 10 datasets with 10 random seeds. More parameter settings on data simulation and the simulation process are in Methods.

The software versions and parameters used for the baseline algorithms are in Methods. For LinRace, the major hyperparameters are *λ*_1_ and *λ*_2_, which are weights for different terms in the likelihood function (Method). we use the default settings, *λ*_1_ = 10 and *λ*_2_ = 1, for all datasets. Although we provide default values for *λ*_1_ and *λ*_2_, we show that LinRace is not sensitive to changes in these parameters. In Supplementary Fig. 2, we show the performances of LinRace are similar under various parameter settings for *λ*_1_ and *λ*_2_. Other parameters used for LinRace are specified in Methods.

As the states of cells and the cell state tree inferred in Fig. 1 Step B are used to infer the states of ancestral cells and calculate the likelihood (Fig. 1 Step C), the accuracy of the inferred cell states and cell state tree can affect the lineage tree reconstruction accuracy of LinRace. With the ground truth cell states and cell state tree provided by TedSim, we can investigate the effect of cell state and cell state tree inference on the final performance of LinRace. To do this, we run LinRace in two modes: (1) using the ground truth cell states and the cell state tree, this mode is denoted as LinRace-TST (True State Tree); (2) using inferred cell states and the cell state tree, and this mode is denoted as LinRace-IST (Inferred State Tree). We use Slingshot [34] to infer cell states and the cell state tree from the gene expression data. As Slingshot does not infer the direction of trajectories, it is a common practice to assign a root cell so that we obtain a directed trajectory. We randomly selected a cell from the root cell state in the true cell state tree to provide this cell to Slingshot as the root cell.

Evaluating the predicted cell division trees involves calculating the distance or similarity between the predicted and ground truth trees, which is not a trivial task due to the large variety of tree structures. Aiming to provide a comprehensive evaluation, we use three different metrics, RF (Robinson-Foulds) distance [26], Nye Similarity [22] and Clustering Info Distance (CID) [32] (Methods). For RF distance and CID, lower is better and for Nye Similarity, higher is better. Settings of data simulation and parameters for running all the methods are in Supplementary Note 2 and Methods.

The results are shown in Fig. 2 (on datasets with dropouts) and Supplementary Fig. 3 (on datasets without dropouts). The results of LinRace-TST, LinRace-IST and baseline methods on all simulated datasets with 1024 cells are shown in Fig. 2a-c. Nye similarity and CID failed to run on the datasets with 4096 cells, so we present the RF distance for datasets with 4096 cells (Fig. 2d, Supplementary Fig. 3d). First, we observe that the two modes of LinRace, LinRace-TST and LinRace-IST, clearly outperform all other methods using all three metrics on datasets with 1024 cells (Fig. 2a-c) and the RF distance on datasets with 4096 cells (Fig. 2d). Comparing Fig. 2a and Fig. 2d, the improvement of LinRace over other methods is even larger with 4096 cells, confirming its effectiveness on large datasets. Second, most methods show a similar trend when the mutation rate changes, except for LinTIMaT. Comparing results on data with dropouts (Fig. 2) and without dropouts (Supplementary Fig. 3), most methods gain much better accuracy on data without dropouts, except for LinTIMaT. Overall, LinTIMaT is not sensitive to the quality of the barcode data, however, its performance is consistently worse than almost all other methods including those using only barcode data, which indicates that it does not take full advantage of the barcode data. Furthermore, it uses a whole-tree local search strategy, which limits its effectiveness on large trees with 1024 leaves.

**Fig. 2.**
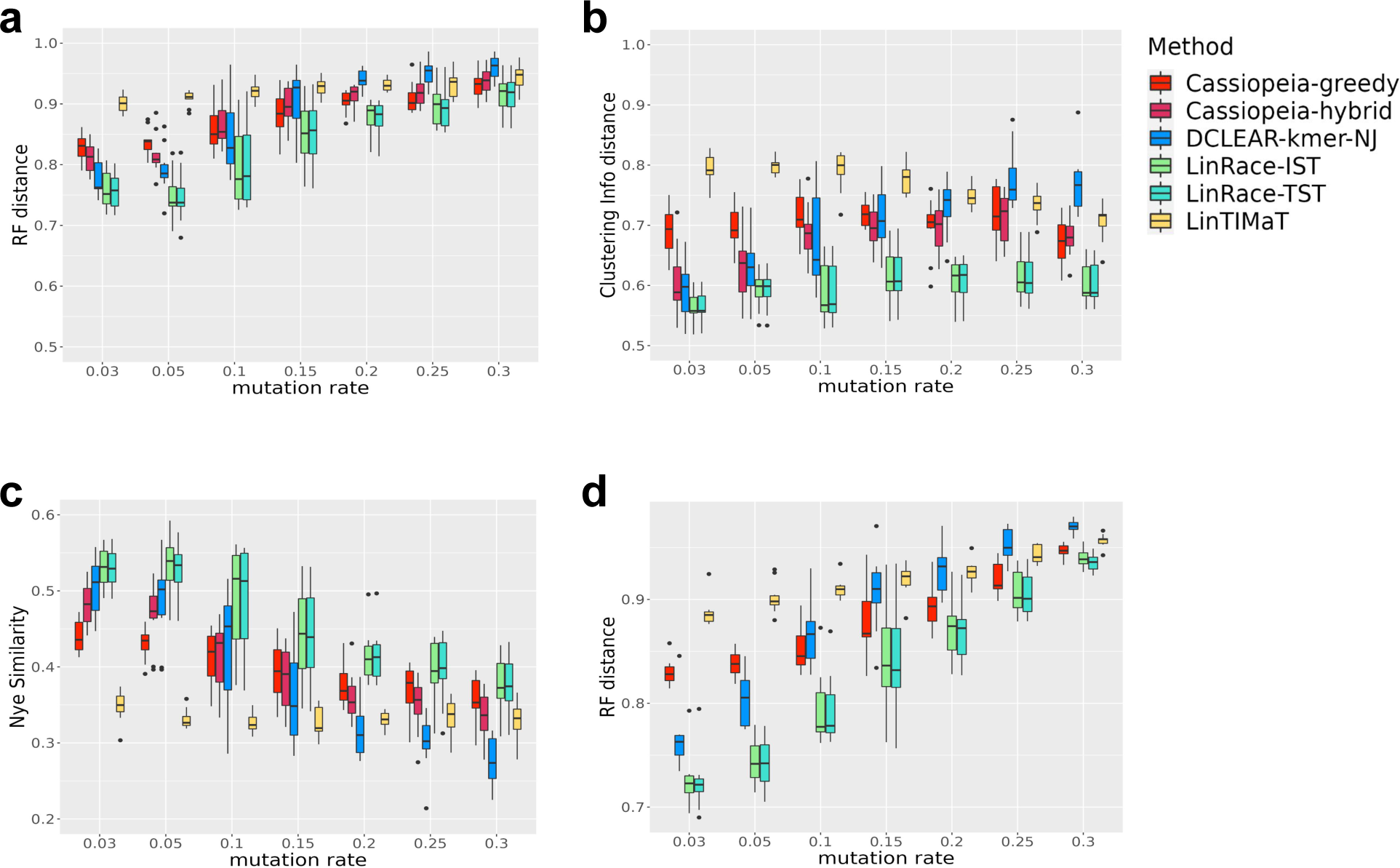
Benchmarking results for lineage reconstruction methods on TedSim simulated datasets with dropouts. **a-c** Comparisons of LinRace (LinRace-IST and LinRace-TST) and other methods on TedSim simulated datasets using RF distance, CID and Nye similarity on datasets with 1024 cells. RF distance, Nye similarity and CID all have the range of [0, 1]. For both RF distance and CID, lower is better, and for Nye similarity, higher values indicate better performance. The detailed descriptions for simulation and method settings can be found in Methods. **d** Comparisons of LinRace (LinRace-IST and LinRace-TST) and other methods on TedSim simulated datasets using RF distance on datasets with 4096 cells.

The trends of the performance of all methods except for LinTIMaT with the increase of mutation rate are expected and are consistent with the observation in [29]. For these methods, the optimal mutation rate for tree reconstruction is in the range of 0.03-0.05 with dropouts (Fig. 2). Larger mutation rates cause more excision dropouts to occur, thus the quality of barcode data decreases, therefore, the performances decrease correspondingly. To quantify the quality of barcode data, we design a score named *reconstruction potential* (Methods). When calculating this, we make use of the known barcodes of ancestral cells from simulation, and count how many edges that connect a parent cell with a daughter cell have at least one mutation introduced from the parent to the daughter. If there is at least one mutation on this edge, it means it is possible to reconstruct this edge of the tree (though it can still be very hard), which corresponds to a split of all leaves cells. If there is no mutation on the edge, we consider that the split can not be reconstructed. Supplementary Fig. 3e shows the reconstruction potential of all datasets with 1024 cells with the change of mutation rates (blue boxes). The trends of the reconstruction potential with the mutation rates confirm the decreasing data quality as mutation rates become too large. Red boxes in Supplementary Fig. 3e show the RF distances of LinRace results. RF distance can be compared with reconstruction potential as they are both based on the ratio of correctly reconstructed splits (Methods). The RF distance of LinRace can sometimes (*µ* = 0.1 without dropouts) exceed the reconstruction potential, which can be attributed to refined local structures inferred from the gene expression data.

LinRace-TST and LinRace-IST, overall show similar performances (Fig. 2). This similarity in performance indicates that LinRace is robust to inference errors in the cell states and cell state tree, which confirms the applicability of LinRace on real datasets, where dropouts are present and true cell states and cell state trees are not available. Since Slingshot was run on each dataset obtained with a given parameter setting and random seed separately, the inferred cell state tree for each dataset can be different, as visualized in Supplementary Fig. 4b. Additionally, when inferring cell states, the ground truth number of cell states is usually unknown. In our evaluation, we used a generic parameter of 7 for the number of states for both simulated and the *C. elegans* real data (Methods). However, it should be noted that the ground truth number of cell states for the simulated data set is 52 (Supplementary Fig. 4a). Despite this difference, our results demonstrate that LinRace is robust to the number of cell states, as evidenced by the comparable performances of LinRace-TST and LinRace-IST.

In the scenario where the barcode data quality is ideal, that is, with no dropouts and with optimal mutation rate (from 0.1 to 0.2 without dropouts), DCLEAR-kmer, which uses only barcode data, can achieve comparable (Supplementary Fig. 3) performance as LinRace, which means leveraging the gene expression data does not yield to significant improvement in this particular case. However, it is important to emphasize that the current lineage barcode data available is far from achieving such ideal quality. Therefore, incorporating gene expression data in a suitable manner becomes crucial for accurate lineage reconstruction.

#### Computational efficiency analysis

The computational efficiency of tree reconstruction algorithms is an important aspect to consider. The computational cost of calculating the likelihood as well as the size of the tree space increases super-exponentially with the number of leaf cells. Therefore, most existing methods such as DCLEAR and Cassiopeia use greedy heuristics to enable efficient tree reconstruction.

In LinRace, we perform local search to learn subtrees on the cells with exactly the same barcode, therefore, for each dataset, the size of the subtrees LinRace needs to learn can vary for every unique barcode. The number of iterations is one of the key parameters for local search algorithms, and often, more iterations are needed for larger trees. We adopt a dynamic manner to set the number of iterations for each subtree based on the Catalan number [3] (Methods), such that smaller subtrees use fewer iterations than larger subtrees, to improve the efficiency of the algorithm.

Supplementary Fig. 5 shows the comparison of LinRace with other state-of-the-art methods in terms of running time on different numbers of input cells. LinRace runs much faster than LinTIMaT, which is the other integrated method that uses both barcode and gene expression data. Compared to barcode-only tree reconstruction methods, Cassiopeia-greedy and DCLEAR, LinRace has a similar running time when the number of cells is small, and its running time grows faster than these two methods when the number of cells increases, in order to search for subtrees that maximize the likelihood function. LinTIMaT has the largest running time despite implementation in Java.

### 2.3 Evaluating LinRace and baseline methods on real *C. elegans* dataset

The ideal scenario to evaluate lineage tree reconstruction methods is to have the following information: the gene expression data and barcode data for single cells, and the ground truth cell lineage tree. While it is rare for experimental data to have ground truth cell lineage trees, *Caenorhabditis elegans* (*C. elegans*) is one of the few species that have the exact cell lineage resolved [23] with single-cell gene expression data measured. To obtain paired single-cell gene expression and barcode data for this system, we adopt a strategy used in [38] to simulate the barcode data from the known lineage tree. Therefore, we combine the measured gene expression data of *C. elegans* and simulated lineage barcode data from TedSim using the true lineage tree. We then apply different lineage reconstruction methods and compare the reconstructed lineage trees with the ground truth tree. When simulating the barcode data, we again vary the mutation rate and the existence of dropouts.

The dataset is obtained from Liu *et al.* [19], who profiled the gene expression lineage of 93 genes in 363 specific cells from L1 stage larvae. They used knowledge of the cell number, morphology of the cell nuclei, and their relative position with respect to each other to develop an automatic method to identify specific cells in confocal images of worms expressing a fluorescent reporter, and then measure expression in specific cell nuclei. The Newick format of the true lineage of the L1 larvae is obtained from CeLaVi [29] by Salvador-Martinez *et al.* which is then trimmed to the profiled 363 cells in the dataset (see Fig. 3a). From the visualization of reduced dimensions in Fig. 3b, we can see that the inferred trajectory is able to connect the cell states and forms a continuous manifold. The inferred cell state tree (shown in Supplementary Note 2) from Slingshot [34] is used to calculate the state transition likelihood in LinRace.

**Fig. 3.**
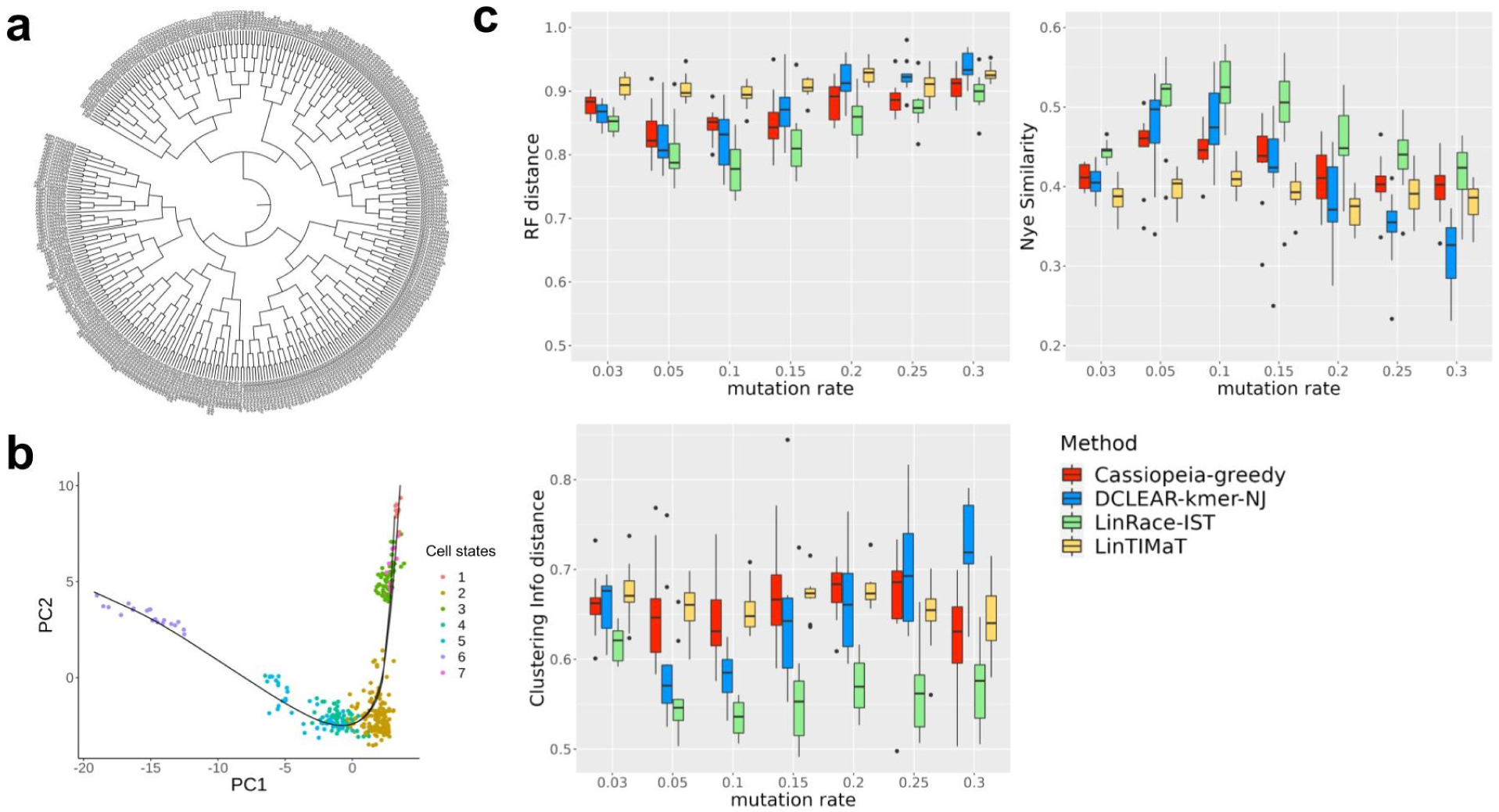
Results of tree reconstruction methods on real *C. elegans* datasets. **a** The ground truth lineage of *C. elegans* at L1 larvae stage. There are 363 profiled cells on the tree while the original L1 larvae lineage has 668 cells. The tree pruning is performed using the ape package in R and the clonal relations between cells are preserved. The tree is visualized using iTOL [18]. **b** 2-D PCA visualization of the gene expression of the *C. elegans* data. After PCA, the first 20 PCs are used for K-means clustering and Slingshot to determine cell states and state trajectories (cell state tree). A detailed description of the processing steps for the gene expression data can be found in Supplementary Note 2. **c** Evaluating result on the *C. elegans* dataset with simulated barcodes. The methods are tested for varying mutation rates with dropout effects. Three metrics, RF distance (lower is better), Nye similarity (higher is better), and CID (lower is better) are presented.

From the results in Fig. 3c and Supplementary Fig. 6, we can see that LinRace outperforms the state-of-the-art methods consistently for varying mutation rates, with (Fig. 3c) or without (Supplementary Fig. 6) dropouts. The overall results are consistent with the evaluation results on simulated datasets. These results not only show the advantages of LinRace over baseline methods on real data but also confirm that LinRace does not need a true cell state tree to be effective. As long as the trajectory inference method captures the relative local relations between cell states, our likelihood function can effectively evaluate a candidate lineage tree based on the raw expression and state transitions. Comparing the performances of the methods with and without dropouts, we can see that LinRace outperforms other methods even more on datasets with dropouts, which indicates that integrated methods like LinRace are able to utilize the gene expression data to compensate for the loss of information caused by dropouts in barcode data. On the other hand, despite using gene expression data, LinTIMaT does not perform better than the two barcode-based methods with dropouts, DCLEAR and Cassiopeia. The reason can be two-fold: first, LinTIMaT runs local search on the whole lineage tree which allows it to explore only a small proportion of the search space, as optimizing this tree as a whole is an NP-hard problem [5]; second, the design of their likelihood function is based on an over-simplified assumption on the relationship between gene expression and barcode data.

### 2.4 LinRace reveals ancestral state transitions of zebrafish brain cells

In most studies that present jointly profiled scRNA-seq and lineage barcode data [1,2,4,13,20,25,31,33], the barcode data and the gene expression data are processed and analyzed separately, where the barcode data is used for building the cell lineage and the gene expression data is used to infer the cell types (Supplementary Fig. 7a). Due to the poor quality of the barcode data, the lineage tree tends to have a relatively low resolution which is reflected by the shallow depth and the small number of internal nodes. Hundreds of cells of different cell types can be connected to the same node and their relative clonal relationships are unknown.

In this section, we show that LinRace can be used to obtain cell lineage trees with better local resolution than the state-of-the-art methods, as well as the cell states of ancestral cells, which is a unique function of LinRace. The finer local structure of the inferred lineage tree inferred by LinRace together with the cell states allows us to identify the location of symmetric and asymmetric divisions in the reconstructed lineage and obtain a picture of how cell types are formed during cell divisions.

We used a zebrafish brain dataset from scGESTAULT [25] and reconstructed the cell lineage using LinRace and other baseline methods. When running LinRace, we used inferred cell states from *k*-means and inferred cell state tree from Slingshot (Supplementary Fig. 7b). The reconstructed cell division trees are visualized in Fig. 4a and Supplementary Fig. 8. We are not able to provide quantitative accuracy of the reconstructed trees as there does not exist a ground true lineage tree. Instead, we compare the resolutions of the trees using the depth and number of internal nodes of the reconstructed trees. From Fig. 4b, we see that the LinRace reconstructed lineage tree has more depth and inferred internal nodes than the Cassiopeia tree and LinTIMaT tree. For DCLEAR which also infers a bifurcating tree, the local splits between cells with the same barcode are randomly decided which does not provide any biological insights, and its number of internal nodes is almost the same as that of LinRace but its depth is much higher means that the DCLEAR tree is much less balanced than the LinRace tree (Supplementary Fig. 8).

**Fig. 4.**
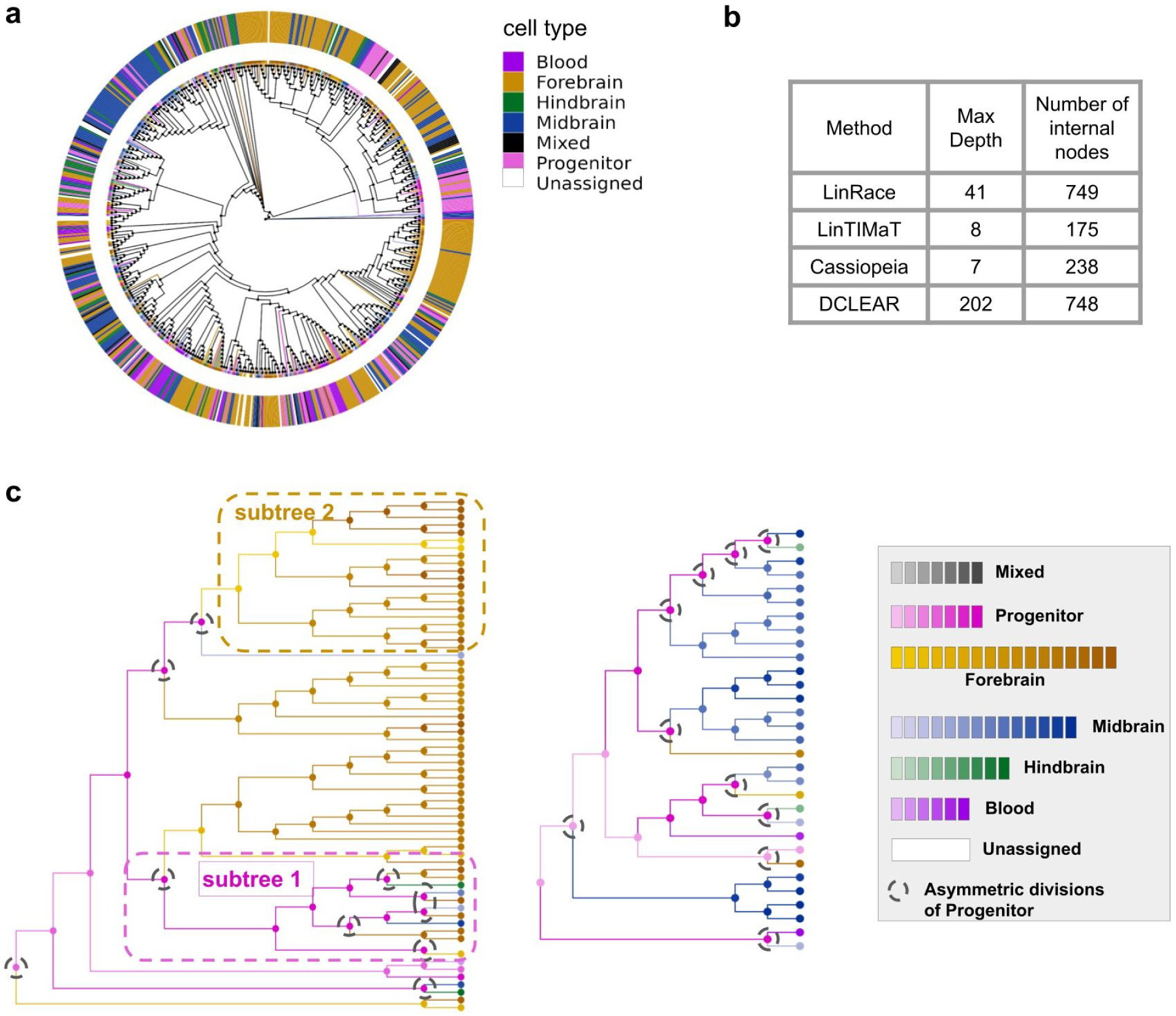
Reconstructed trees of ZF1-F3 sample (750 cells) from scGESTAULT datasets in [25]. **a** LinRace reconstructed tree. The outer ring represents major cell type assignments and the inner colors on the edges show detailed intermediate cell type assignments. **b** Properties of the reconstructed trees from different methods. The max depth means the maximum total edge length going from the root to any leaf cell. **c** A detailed look of two GES (Gene Expression Subtrees) in LinRace reconstructed tree with inferred ancestral states. In the left GES, Subtree 1 shows a subtree of progenitor’s self-renewal and subtree 2 shows a subtree of differentiated Forebrain population. Black dashed circles denote asymmetric divisions of progenitor cells. The annotated leaf cell states are from the original data paper, and the ancestral states of all hidden nodes are inferred using LinRace. Detailed descriptions of the processing steps for the gene expression data are in Supplementary Note 2.

Fig. 4c shows two Gene Expression Subtrees (GES) reconstructed by LinRace where the color of nodes (including both leaf nodes and ancestral nodes) represent the cell types of cells, as annotated in the original paper [25]. Although LinRace used inferred cell states and cell state tree when performing tree reconstruction, we analyze the reconstructed lineages by labeling the cells with annotated cell types. To obtain the states of ancestral cells, we ran the ancestral state inference procedure (Methods) again using the annotated cell types of leaf cells and a cell state tree of these cell types (Supplementary Note 2). After obtaining the cell states of both leaf and ancestral cells, we can examine the cell state changes on the reconstructed lineages, and observe how the progenitor cells adopt asymmetric divisions to generate different clones of various terminal cell types. The two GESs are dominated by forebrain and midbrain cell types, respectively (Fig. 4c left and middle). In each GES, the cells have the same barcode, so without using gene expression information, no meaningful structure can be reconstructed for each GES. LinRace reconstructs these subtrees along with the states of ancestral cells. From Fig. 4c, we can see where some progenitors undergo asymmetric divisions and differentiate into various cell type after self-renewal (Subtree 1 in Fig. 4c left), while cells that are have entered a terminal cell type (forebrain) divide mostly symmetrically to maintain the sustainable activity of neurogenesis (Subtree 2 in Fig. 4c left). In Fig. 4c middle, multiple symmetric divisions happen to lead to the Midbrain cell types.

Finally, from the visualization of reconstructed cell lineage trees with different methods (Fig. 4a and Supplementary Fig. 8), we can see that compared to the other methods, LinRace is able to reconstruct a lineage tree with a realistic distribution of cell types, where while cells with the same cell types tend to be located together in the reconstructed lineage tree, the same cell type can appear in different subtrees, shows the “partial consistency between transcriptome similarity and barcode similarity” in real data. In the next section, we further show this phenomenon using another dataset of the mouse embryo. Further, we show how asymmetric divisions explain this phenomenon and the conjecture of varying cell differentiation speeds on different lineages.

### 2.5 LinRace helps to answer the sources of diverse cell types in the mouse embryo data

We applied LinRace to an early mouse embryo dataset from Chan *et al* [4] to infer the cell division tree. The mouse embryo dataset contains 9707 cells of 34 annotated cell types from the authors (Fig. 5a). On this dataset, we not only analyze how the cell types are distributed over the cell division tree but also ask if any lineage signature exists in a cell’s gene expression profile.

**Fig. 5.**
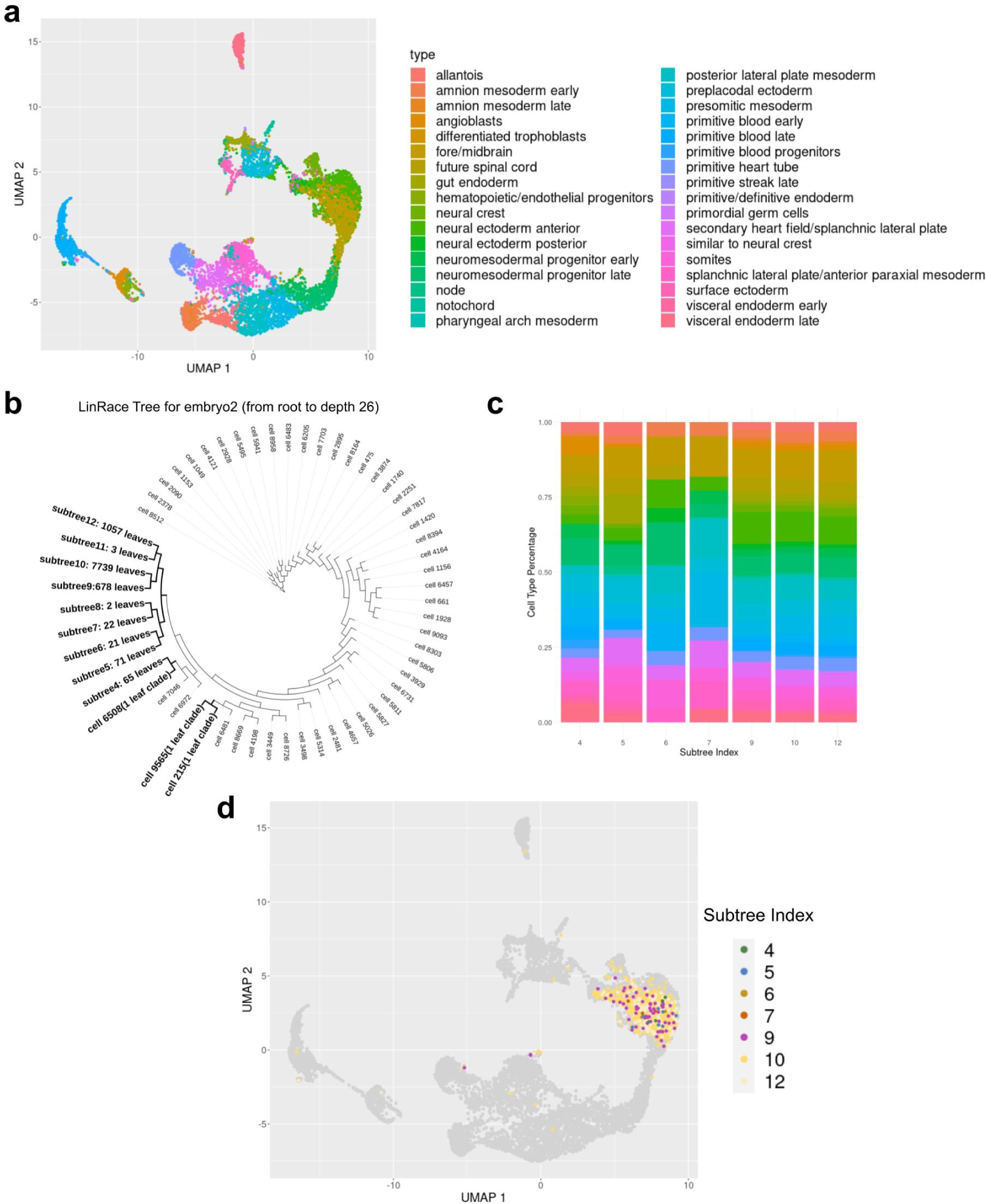
Visualizing inconsistencies between cell states and lineages. **a** UMAP visualization of the early mouse embryo dataset with cell type annotation from [4]. **b** LinRace reconstructed tree of the dataset. Only branches from root to *depth* = 26 is shown here, where the remaining subtrees are shown as leaf nodes, with the numbers of leaf cells attached to them. Three one-leaf clades are included which means they are leaf nodes at *depth* = 26. **c** Cell type compositions under different subtrees. The subtrees are obtained by cutting the whole lineage at the same depth (total edge length from the root to the cutting point.) 12 subtrees are obtained by cutting the lineage tree at *depth* = 26 and only subtrees have more than 10 leaves are shown here. The y-axis shows the percentage of each cell type in a under each subtree. **d** UMAP visualization of the Fore/Midbrain cell type by the clonal IDs. Only subtrees have more than 20 leaves are labeled and cells that do not belong to these subtrees are colored black and labeled as “0”.

We first cut the reconstructed tree at depth 26 and obtained subtrees with their roots at a distance of 26 to the root of the complete tree (Fig. 5b). First, we investigate the cell type composition of the subtrees. The cell type composition of 7 subtrees with at least 10 leaves is shown in Fig. 5c. Each subtree consists of multiple cell types (colors are consistent with the color legend of Fig. 5a), and the same cell type appears in different subtrees, again showing the “partial consistency between transcriptome similarity and barcode similarity”.

Next, we take the cells that belong to the same cell type (“Fore/Midbrain” in Fig. 5a) and look for differences that potentially exist between cells from different lineages but with the same cell type. We visualize the Fore/Midbrain cells labeled by their clonal IDs (same as the subtree IDs, Fig. 5d). In the UMAP space, cells from different clones are mixed and do not show any cluster pattern. Differential expression (DE) analysis between cells in different subtrees and the same cell type did not show DE genes whose functions can clearly cause clonal differences. These observations indicate that there are no significant lineage-affiliated features in the reduced dimensions of the transcriptomic data, which is in line with the hypothesis that the gene expressions of mature cell types are dominated by their functionality, not their lineage identity, as previously reported in [23].

Further, it is a common observation that in a scRNA-seq dataset, although all the cells are profiled at the same time, the cells can have different pseudotime representing varying stages of development or distinct cell states. In the context of the cell division tree that gives rise to the leaf cells, we conjecture that leaf cells at a later stage of development originate from lineages with fast differentiation speed, potentially involving frequent asymmetric divisions, and leaf cells at an early stage of development can originate from lineages with slow differentiation speed.

## 3 Discussion

In this paper, we present LinRace, an integrated method that combines the lineage barcode and gene expression using the asymmetric cell division model to reconstruct cell division trees. Compared to the state-of-the-art methods, LinRace has the following advantages: (1) Using the combined framework of NJ and maximum likelihood, LinRace outperforms the state-of-the-art methods on both simulated and real datasets; (2) LinRace proposes a novel likelihood function that takes into account mRNA counts, cell states and state transitions to find the lineage tree that best explains the observed gene expression data; (3) LinRace is able to infer the cell states of ancestral and provides insights on the process of generating new cell types; (4) LinRace performs a local search on subtrees using a dynamic number of iterations based on the size of the subtrees which makes it computationally efficient compared to other tree search algorithms.

Reconstructing the cell division tree from lineage tracing barcodes is similar to the problem of inferring phylogenetic trees from genome data in the field of evolutionary biology [14]. However, the cell division tree reconstruction is even more challenging due to the much larger number of leaves (single cells) and the short lineage tracing barcodes with dropouts. Incorporating the single-cell gene expression data is a promising direction, but developing such integrated methods requires modeling the relationships between the two modalities, the lineage barcode, and the gene expression. LinRace adopts the asymmetric division model, which explains phenomena in real data, increases reconstruction accuracy, and allows for the estimation of ancestral cell states.

LinRace assumes that cell state transitions are irreversible, which is generally true under homeostatic conditions. However, reversible state transitions are also known to exist. To incorporate reversible transitions, it may be necessary to expand the definition of the cell state tree to a cell state network in future work. In addition to developing new computational methods, it is also important that the data quality is improved with advances in technology, in order to build the tree of cell division and cell differentiation with high accuracy.

## 4 Methods

### 4.1 The asymmetric division model accounts for cell state changes in the lineage tree

The asymmetric division of cells is a key process that leads to multiple cell types and different cell differentiation speeds on different clones on the cell lineage tree [6,21]. We show that the asymmetric division model can account for various scenarios of cell state change on a cell division tree reviewed in [36]. Moreover, it was shown to lead to realistic paired lineage barcode and gene expression data [24]. An asymmetric division results in two unequal daughter cells from the parent cell, one of which differentiates into a natural “next-state” (according to the cell state tree) while the other remains in the same cell state as the parent.

Wagner and Klein [36] reviewed hypothetical scenarios of restricted lineage trajectories unfolding on a state manifold of gene expressions. In the review, state convergence represents two or more distinct fate trajectories converging onto the same final position on a state manifold, and state divergence represents the reverse process where one trajectory bifurcates into two or more distinct fate trajectories. These seemingly contradictory scenarios can happen due to asymmetric cell divisions. Asymmetric divisions can cause one cell state to generate two distinct cell states, resulting in state divergence; and cells on distant lineage trees can also divide asymmetrically into the same cell state, resulting in state convergence. In LinRace, we account for both symmetric and asymmetric cell divisions in our likelihood function using the asymmetric division likelihood which estimates the prior probability of asymmetric and symmetric divisions based on the mutated states in the observed cells. We assume that state transitions on all parent-child cell edges are independent, and also consider the stochasticity of cells’ developmental speeds by varying the number of states traversed for each state transition. That the symmetric and asymmetric cell divisions happen on the cell lineage tree according to certain probabilities are called the asymmetric division model in this paper. In LinRace, the asymmetric division model is also used in inferring ancestral cell states.

### 4.2 Reconstructing lineage backbone from the lineage barcode data

The lineage barcode of a cell is represented as a character vector of length equal to the number of target sites as designed by the CRISPR/Cas9 lineage recorder. Each character represents a state of the target site, which can be a mutated state, an unmutated state, or a dropout state. We use “0” to represent the unmutated state, and each unique non-zero character represents a unique mutation state, regardless of the position where the mutation is observed. The dropout state is denoted as the “-” character. For the barcode data, we assume that at the root of the cell lineage, an unedited DNA sequence (all unmutated states) is introduced. During cell divisions, unmutated states can potentially mutate and will never mutate again, except the dropout state. Given *N* cells and *M* targets, the lineage barcode data is a *N × M* matrix. For the barcode data, we assume that at the root of the cell lineage, an unedited DNA sequence is introduced. During cell divisions, unmutated target sites can potentially mutate and will never mutate back to the original state (except dropout state).

In LinRace, we utilize a Hamming distance-based NJ method to infer the “backbone lineage tree”. Since multiple cells can have the same barcode in the dataset, running Neighbor Joining on the *N × M* matrix will result in merging cells with the same barcode in some random order. With a given lineage barcode matrix, we first transform it into a *K × M* matrix where each row represents a unique barcode in the original data, and then Neighbor Joining is applied to the *K × M* matrix to get the lineage tree of unique barcodes (Fig. 1). We call this tree the reconstructed lineage backbone, or tree backbone (denoted by *T*_0_).

### 4.3 Inferring cell states for ancestral cells

To calculate the likelihood of a candidate tree based on the gene expression profiles of the leaves, we first need to infer the states of cells at ancestral nodes using the states of leaf nodes and the cell state tree that are inferred during Fig. 1 Step B. The inference of ancestral cell state follows the rule of asymmetric division. From leaves to the root of the candidate lineage tree, we consider the following cases: (1) If the daughter cells have the same cell state, their parent is assigned the same state, meaning the parent cell divides symmetrically. (2) If the daughter cells have different cell states, the parent cell will have the Most Recent Common Ancestor (MRCA) cell state based on the cell state tree. The ancestral cell states in trees in Fig. 1 Step C follow these two rules given the cell state tree learned from Step B. This ancestral state inference process allows one cell to divide into two cells with different cell states. If two daughter cell states belong to the same differentiation path (a path from the root state to a leaf state), the parent cell will be in the same cell state as the earlier cell state between the two.

### 4.4 Finding subtree topology using a maximum likelihood method

Cells with identical barcode form a star tree in the initial NJ tree *T*_0_ (See Fig. 1). LinRace utilizes the gene expression data to learn the bifurcating tree topology of these cells. The learned bifurcating trees, termed Gene Expression Subtrees (GES), are then attached to the tree backbone to yield the full cell lineage tree. We design a maximum likelihood scoring function and local search strategy to find the GES.

#### Likelihood of a candidate lineage tree

We design a likelihood function to evaluate how well a candidate lineage tree explains the observed gene expression data. We use the ancestral state inference step mentioned above to determine the cell states of all nodes on the tree. Then, we can calculate the likelihood of the tree, which consists of three terms: *state transition likelihood*, *asymmetric division likelihood*, and *neighbor distance likelihood*. The first two terms use the cell state information of cells, and the last uses the gene expression profiles of the cells.

The *state transition likelihood* represents the probability of transitions between cell states on the edges of the lineage tree. We adopt the assumption that the state transition on each edge is independent of other transition events (this assumption is commonly used in phylogenetic tree reconstruction) so that we can write the state transition likelihood of a given lineage tree *T* = (*E, V*) as:

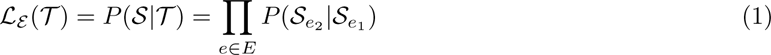

where *e* = (*e*_1_*, e*_2_) represents an edge on the tree graph from node *e*_1_ to *e*_2_, *S* denotes the cell state assignments of all cells, and *S_e_*_1_and *S_e_*_2_ denotes the cell states of the two cells *e*_1_ and *e*_2_ connected by *e* ∈ *E*.

The state transitions of cells’ gene expressions are governed by an underlying developmental cell state tree, which is inferred from the gene expression data using Slingshot (Fig. 1 Step B). The cell state tree guides the cells to differentiate into certain future cell states irreversibly. For any two states on the cell state tree, there can exist at most one path(a sequence of connected, directed edges) that links these two states. Therefore, when the states of a pair of ancestor-descendant, denoted as (*S_e_*_1_*, S_e_*_2_) between two cells (*e*_1_*, e*_2_), the transfer probability *P*(*S_e_*_2_*|S_e_*_1_) is calculated as follows:

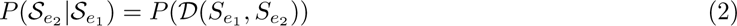

where *D*(*S_e_*_1_*, S_e_*_2_) represents the graph geodesic from *S_e_*_1_ to *S_e_*_2_ on the cell state tree. If *D*(*S_e_*_1_*, S_e_*_2_) = + inf, a penalty of *−*50 is applied to the log likelihood. The probability for every distinct state transition is given as follows:

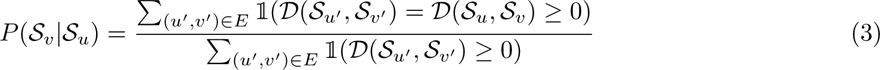

The *asymmetric division likelihood* considers the asymmetric divisions in the lineage tree and it is defined as follows:

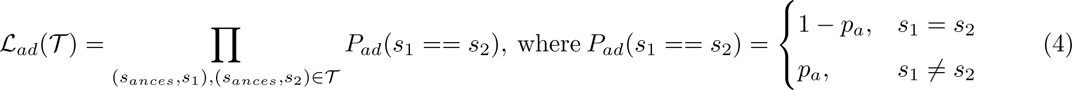

where *s_ances_* is the state of a parent cell and *s*_1_*, s*_2_ are the states of the two descendant cells. The asymmetric division rate *p_a_* can be determined using prior knowledge or inferred from the fraction of observed asymmetric neighbors in the lineage tree.

For *neighbor distance likelihood*, we look at cells that are siblings (having the same parent node) on the candidate lineage tree, and use the transition probability from diffusion map [11] to evaluate if the two cells are locally connected on the developmental trajectories of the gene expression data. Even though in general when asymmetric division happens, the two daughter cells do not have the same cell states thus their transcriptomes are not very similar, we consider that many cells at the leaves of the lineage tree are terminal state cells, and asymmetric divisions are less prominent when more cells are at terminal states. We denote the set of measured cells by *Ω*. First, for all pairs of cells (*x_i_, x_j_*) in *Ω*, we calculate the pairwise distance of cells’ gene expression data using the radial basis function (RBF) kernel 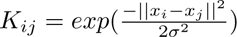. Then, the transition probability between any two cells can be calculated as follows:

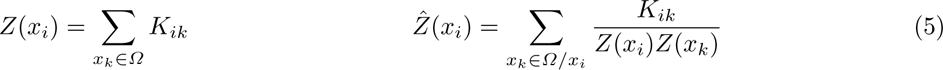

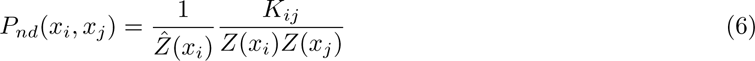

and the neighbor distance likelihood can be calculated as:

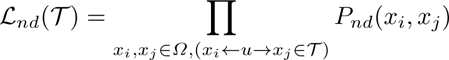

Finally, the total likelihood is calculated as:

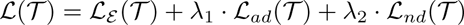

where *λ*_1_ and *λ*_2_ are hyperparameters.

To find the best GES based on our likelihood function, we utilize hill-climbing local search to search in the tree space (Fig. 1 Step C). In order to propose a new tree from a current tree, we adopted random Subtree Swapping (rSS), which is a derivative of the widely used random Subtree Pruning and Regrafting (rSPR) [8]. We randomly select two nodes on the current tree and prune the subtrees attached to the specific nodes. Then, we regraft either one of the pruned subtrees to the location of the other subtree. The advantage of rSS is that with an initialization of a bifurcating tree, this operation will not disrupt the binary property throughout the searching process. This move can result in most of the applicable topological changes of the tree so we use it as the search technique. At every iteration, a new tree is proposed which is one rSS move away from the current tree. Then, we evaluate the new tree with the likelihood function and compare it with the likelihood of the current tree. If the new tree has a higher likelihood, we move from the current tree to the new one. Repeat the process for every iteration until no better tree can be found within a number of iterations, and the current tree is identified as a local optimal tree. We use a random restart technique after a local optimal tree is found, and try to find as many local optimal as possible within the number of maximum iterations. If multiple local optima are found, we return the tree with the highest likelihood as the optimal GES. The pseudocode of the local search process of LinRace is given in Supplementary Note 1.

### 4.5 Simulating synthetic datasets with TedSim

We use simulated datasets from TedSim which generates paired lineage barcode and gene expression data with ground truth information of the lineage tree. Using the simulated datasets, we are able to benchmark LinRace and other tree reconstruction methods by comparing the reconstructed trees with the ground truth. To simulate with TedSim and test the methods’ potential under various conditions. We use a pre-defined cell state tree (Supplementary Fig. 4) and vary key parameters including mutation rate and the presence of dropout to simulate lineage barcodes of different quality. The selected mutation rate range from 0.05 to 0.4 per target per cell division which covers the realistic ranges of mutation rate in real datasets.

One important advantage of TedSim is that TedSim simulates realistic dropout effects that widely occur in real datasets. In TedSim, when two or more mutation happens at the same cell division, the excision dropout will happen, resulting in deletions of the targets in between. These dropouts can cause a significant decrease in barcode diversity. Comparing the distributions of unique barcodes of real dataset [4] (Supplementary Fig. 1), a simulated dataset with dropout, and a simulated dataset without dropout, we can see that TedSim simulation with dropout is able to generate realistic data with similar negative exponential distribution. In the real dataset of 19019 cells and 2788 unique barcodes, and as mentioned in Results, 1929 of the 2788 barcodes uniquely label one cell; 330 of the 2788 barcodes are shared by two cells. In a simulated dataset of 1024 cells with dropout, 88 cells are uniquely barcoded (no other cell shares the same barcode) and 44 barcodes are shared for 2 cells. This means, 82.8% of the cells share the same barcode with at least two other cells, which is similar to the proportion of 86.4% in the embryo2 dataset in the mouse lineage tracing system. Without dropout, the number of uniquely barcoded cells increases to 336 and 136 barcodes are shared for 2 cells. The proportion of cells that share the same barcode with at least two other cells decreases to 40.6%. TedSim-generated datasets with dropouts reflect the distribution of number of cells using the same barcode reflect the distribution in real data (Supplementary Fig. 1).

The cell state tree and cell states used for simulation are in Supplementary Note 2. The step size parameter in TedSim is used to sample the cell state tree to obtain discretized cell states from the tree. The step size in our experiments is set to be 0.5 which yields 52 discrete cell states in the simulated datasets. Other static simulation parameters can also be found in Supplementary Note 2.

In the experiment with the *C. elegans* dataset, TedSim is used to simulate lineage barcodes on the ground truth lineage tree. We also tune the mutation rate from 0.05 to 0.4 per target per cell division and simulate lineage barcode data with and without dropouts. We choose a smaller number of target sites(*N char* = 9) considering the size of the lineage(363 cells).

### 4.6 Evaluation metrics for tree reconstruction

To evaluate the accuracy of reconstructed lineage trees, we use three tree comparison metrics: RF distance, Nye similarity, and CID (Clustering Information Distance) distance. We use the RF.dist() function from the `phangorn` package to calculate the RF distance between reconstructed lineage trees and ground truth trees from simulated datasets. The function returns the ratio of inconsistent splits between the two trees. Therefore, the distance value ranges from 0 and 1, where 0 represents a perfect reconstructed tree and 1 means the worst reconstructed trees where all splits are different from the true lineage. The RF distance is a very strict metric where it only counts as a correct split if only all leaves on the two sides of the splits are consistent for the two trees.

Therefore, we also introduce another metric, Nye Similarity score [22], which can be regarded as an extended version of RF distance. Nye Similarity also compares each split of edge on the two trees, but instead of giving a binary score(0 or 1 for RF distance), it calculates a score based on the similarity between the two splits. Considering each edge *e* in a tree *T*, the score *s*(*e_i_, e_j_*) is for any pair of edges (*e_i_, e_j_*), *e_i_* ∈ *T*_1_ and *e_j_* ∈ *T*_2_, is obtained by the partition of the leaf nodes. Considering the set of all leaf nodes *L*, we have:

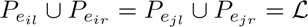

where *P_eil_, P_eir_* are the two disjoint subsets of *L* by the edge *e_i_*, and *P_ejl_, P_ejr_* are the two disjoint subsets by the edge *e_j_*. Then, for *r, s* = *l, r*, the number of leaf nodes shared by the partition can be given as,

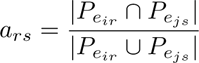

For a pair of splits (*e_i_, e_j_*), the score *s*(*e_i_, e_j_*) is then defined by

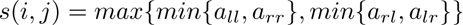

Finally, the Nye Similarity score is derived by maximizing the quantity:

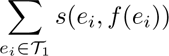

where *f*(*e_i_*) is an alignment of edges for the two trees. The quantity is maximized by finding the best alignment of edges *f*.

Clustering Information Distance (CID) is another information-theoretic generalized RF distance metric. CID calculates a matching score for every split on the tree which is based on Mutual Clustering Information [35]. Such metric, as previously compared [32], has the advantage of being continuous and relatively unbiased towards certain topologies. Using the three metrics, we are able to comprehensively evaluate the tree reconstruction methods, while RF distance provides a basic metric that is also widely used in other papers, the generalized metrics like Nye Similarity and CID provides more detailed and interpretable comparisons.

### 4.7 Running LinRace and baseline algorithms

Here is a summary table of the key packages: lineage inference methods (Cassiopeia, DCLEAR, LinTIMaT), trajectory inference methods (Slingshot), data simulation (TedSim) and evaluation (TreeDist), that are used in this paper:

Major parameters and their settings for LinRace are:

– *λ*_1_ and *λ*_2_, set to be *λ*_1_ = 10 and *λ*_2_ = 1 in all tests.

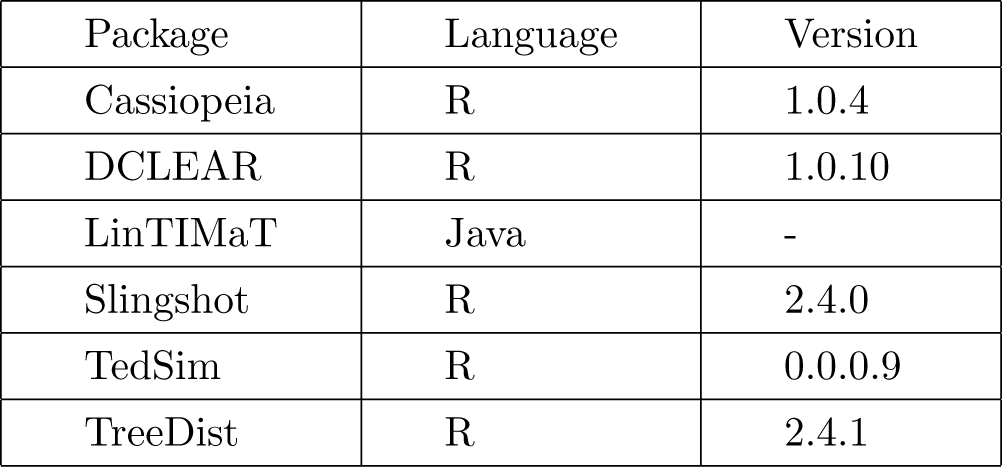

| Package | Language | Version |
| --- | --- | --- |
| Cassiopeia | R | 1.0.4 |
| DCLEAR | R | 1.0.10 |
| LinTIMaT | Java | - |
| Slingshot | R | 2.4.0 |
| TedSim | R | 0.0.0.9 |
| TreeDist | R | 2.4.1 |

– Number of genes kept after filtering: we keep the top 100 highly variable genes for simulated data and …

– Maximum number of iterations for each local search is 500 in all results.

– Asymmetric division rate *p_a_* is set to 0.8 in all tests

Running LinRace-IST also involves the steps of identifying cell states and inferring the cell state tree. *k*-means clustering method is used on the scRNA-seq data to identify cell states and Slingshot is applied to infer the cell state tree, based on the identified cell states. When using simulated data, the number of clusters for *k*-means is set to 7. The same number of clusters is used for the *C. elegans* data. For the scGESTAULT dataset (ZF1-F3), the number of clusters for *k*-means is set to 20. For the mouse embryo data, we used annotated cluster IDs from the paper [4].

As the subtrees which LinRace local search aims to optimize can vary in size, we adopt a dynamic manner to set the number of iterations for each subtree. Denote the *n*^th^ Catalan number [3] by *C*(*n*), then the number of iterations used to optimize a subtree with *n* leaves is: min(*C*(*n*)*, max_<u>i</u>ter*), where *max_<u>i</u>ter* is the maximum number of iterations set as a parameter.

The parameter settings for baseline methods were mostly based on their default parameters. LinTIMaT has three major parameters: the number of genes *gc*, the number of mutation likelihood iterations *mi*, and the number of combined likelihood iterations *ci*. We set *gc* = 100 when running it on simulated data, the zebrafish dataset, and the mouse embryo dataset. For the C. elegans data, we used *gc* = 93 because the dataset has in total 93 genes. *mi* and *ci* were set to 20000 for both simulated and real datasets.

For DCLEAR, we used the “kmer” mode and set the *k−*mer length to 2 for all datasets, which is the default value for this parameter. We also used default values of other parameters for DCLEAR.

For Cassiopeia, Cassiopeia-greedy and Cassiopeia-hybrid are used for 1024 cells. For the Cassiopeia-hybrid, we set the convergence time limit for the ILP solver to be 1000, the maximum potential graph layer size to be 500 and set a cell cutoff between the top greedy solver and the bottom ILP solver to be 20. For the parameter *indel_priors* (which represents the probabilities of particular indels occurring), we have left it empty.

More detailed descriptions of the parameter settings for running the simulation and the lineage reconstruction methods can be found in Supplementary Note 2.

### 4.8 Reconstruction potential of lineage barcode datasets

Given a true lineage tree *T*^∗^ and a character matrix *X* (lineage barcodes) of both observed, leaf cells and hidden, ancestral cells (for TedSim datasets, the lineage barcodes of the ancestral nodes are known), we analyze the *reconstruction potential* which can be compared to the Robinson-Foulds distance of a lineage reconstruction method because both metrics are calculated based on the ratio of the splits on the lineage tree. The idea of the reconstruction potential is: for any given edge that splits all leaf cells into two sets, at least one mutation is required for this split to be reconstructed. Denote the true lineage tree as *T*^∗^ = (*V, E*), where *V* represents the cells on the lineage tree. The reconstruction potential *Q_r_* is given as follows:

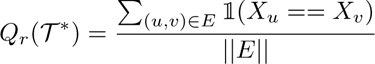

where 1(*x*) is the characteristic function which returns 0 if *x* is TRUE; 0 otherwise. *Q_r_* calculates the fraction of edges that contain at least a mutation.

## 5 DATA AVAILABILITY

LinRace is available at: https://github.com/ZhangLabGT/LinRace.

TedSim is a simulator for paired lineage barcode and gene expression data available at: https://github.com/Galaxeee/TedSim.

Cassiopeia is an end-to-end pipeline for single-cell lineage tracing experiments available at: https://github.com/YosefLab/Cassiopeia.

DCLEAR is an R package for Distance-based Cell LinEAge Reconstruction (DCLEAR) available at: https://github.com/ikwak2/DCLEAR.

LinTIMaT is a statistical method for reconstructing lineages from joint CRISPR-Cas9 mutations and single cell transcriptomic data available at: https://github.com/jessica1338/LinTIMaT.

## Supplementary Information

### 1 Supplementary Figures

**Supplementary Figure 1.**
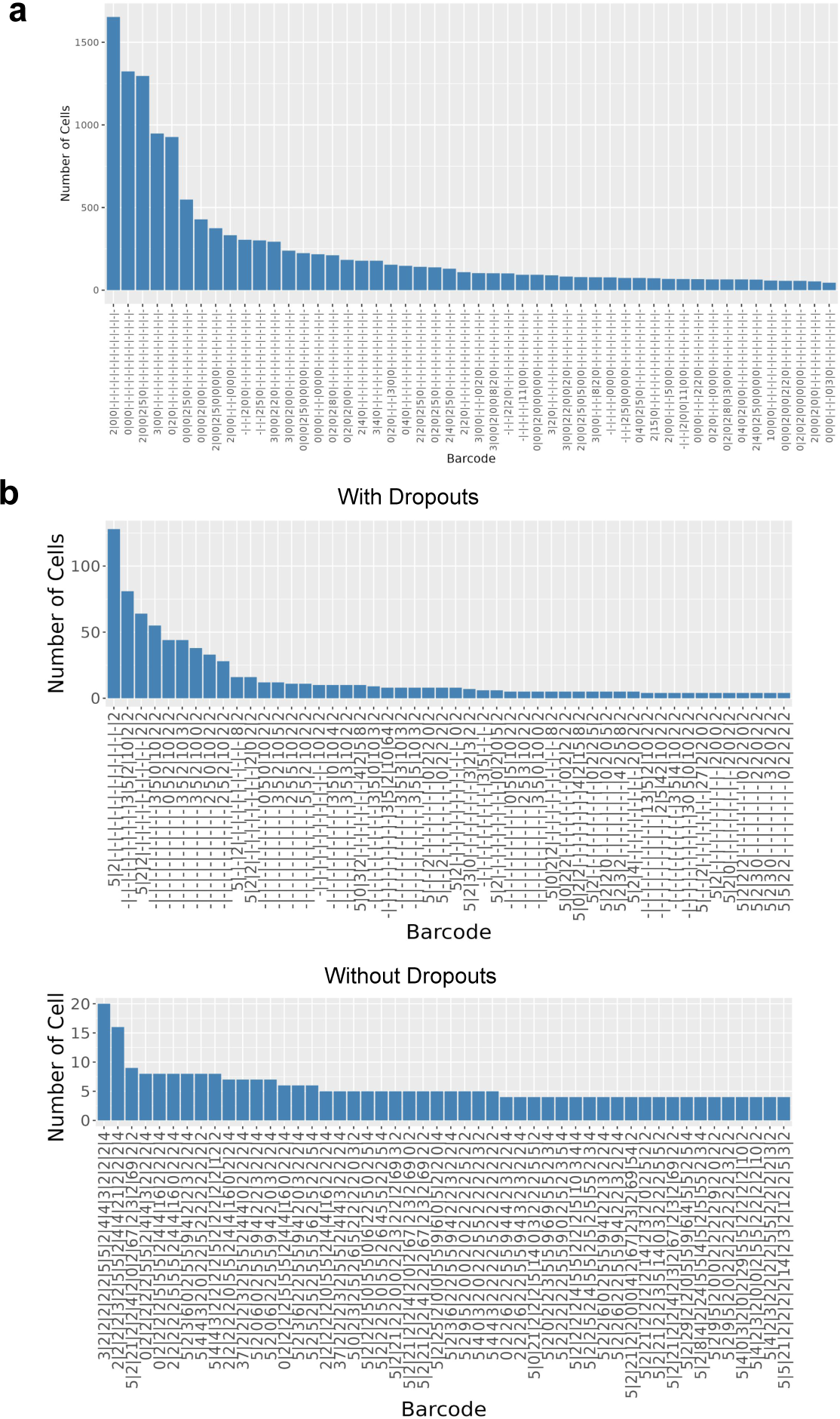
Barcode distributions in both real and TedSim simulated datasets. x-axis shows barcode in the form of character strings, where “0” denotes unmutated state, nonzero numbers denote mutations, and “-” denotes dropouts. Top 50 barcodes with the most cells are selected. y-axis shows the number of cells with the same barcode. **a** Barcode frequencies of the embryo2 dataset in M. Chan *et al*. The dataset has 19019 cells, 18 targets, and a total of 2788 unique barcodes. **b** Barcode frequencies of TedSim simulated datasets, one with dropouts and one without. Both datasets have 1024 cells, 16 targets for the barcode and mutation rate = 0.1.

**Supplementary Figure 2.**
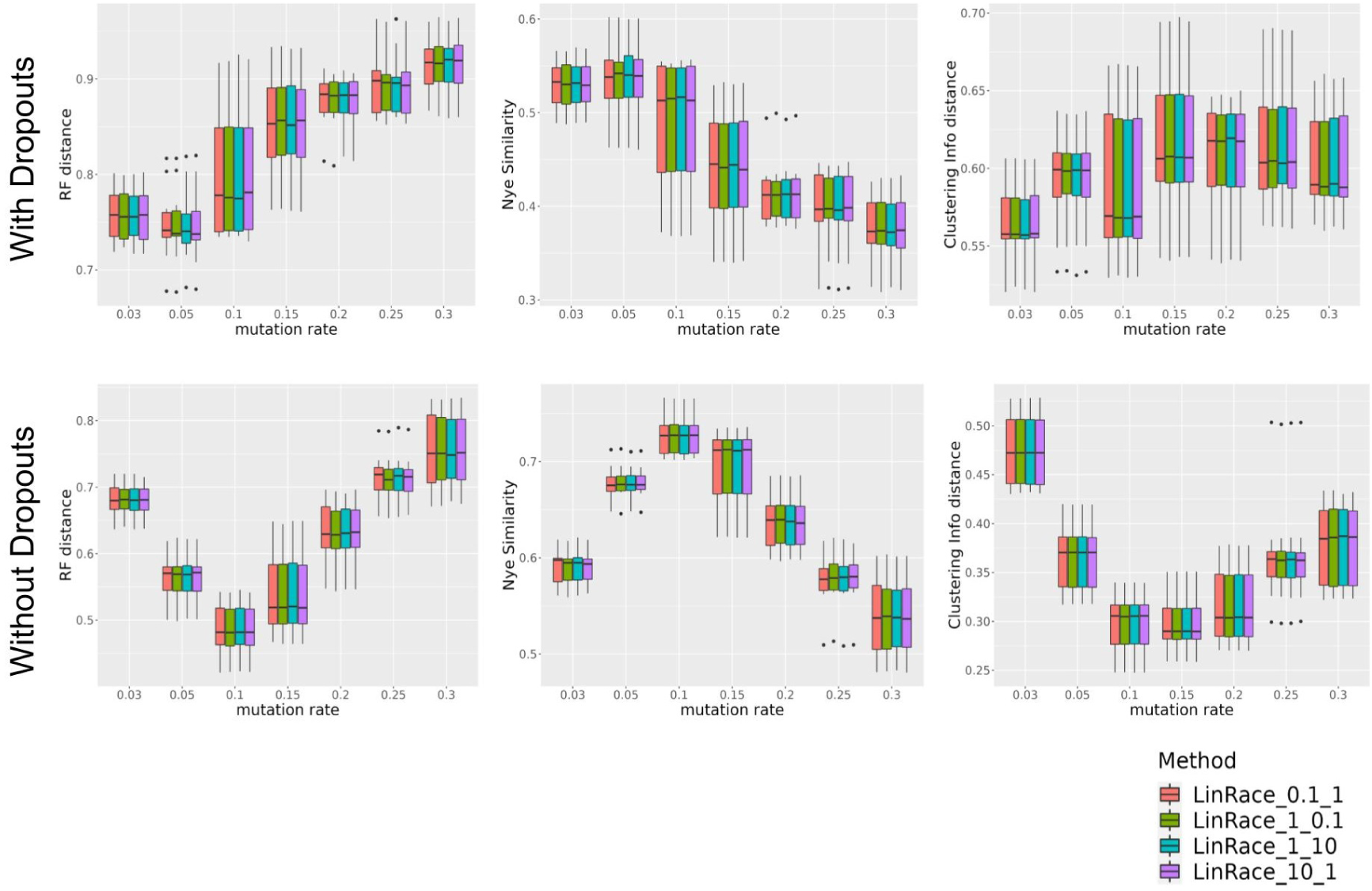
Comparisons of LinRace with different settings of hyperparameters. The first parameter corresponds to *λ*_1_, the weight for asymmetric division likelihood. The second parameter corresponds to *λ*_2_, the weight for neighbor distance likelihood. The datasets used are the same data used for benchmarking the lineage reconstruction methods in Fig. 2a.

**Supplementary Figure 3.**
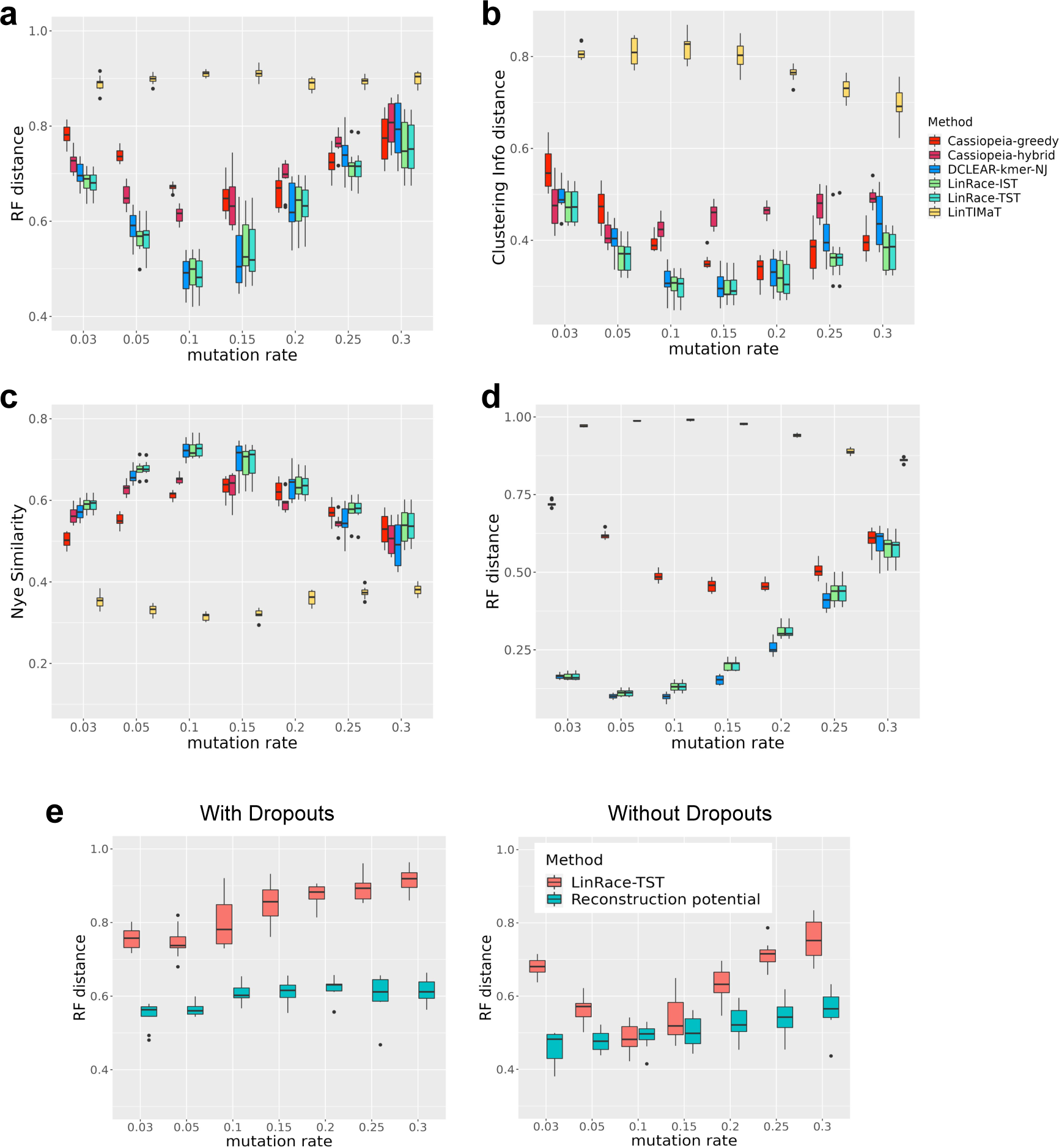
Comparisons of LinRace (LinRace-IST and LinRace-TST) and other methods on TedSim simulated datasets without dropouts. The number of target sites is set to 64. The other simulation settings for gene expression data as well as parameters for running the algorithms are the same as described in Methods and Supplementary Note 2 for 1024 cells. For every combination of parameters, 10 simulated datasets are generated. **a** RF distance on datasets with 1024 cells. **b** Nye similarity on datasets with 1024 cells. **c** CID on datasets with 1024 cells. **d** RF distance on datasets with 4096 cells. Nye similarity and CID all have the range of [0, 1]. For both RF distance and CID, lower is better, and for Nye similarity, higher values indicate better performance. Only RF distance results are given on 4096 cells because Nye similarity and CID are too slow to run on such large datasets. **e** Comparison of the reconstruction potential and LinRace-TST performances. The data used to calculate the reconstruction potential and the RF distance of LinRace-TST reconstructed trees are the same data from Fig. 2 a-c.

**Supplementary Figure 4.**
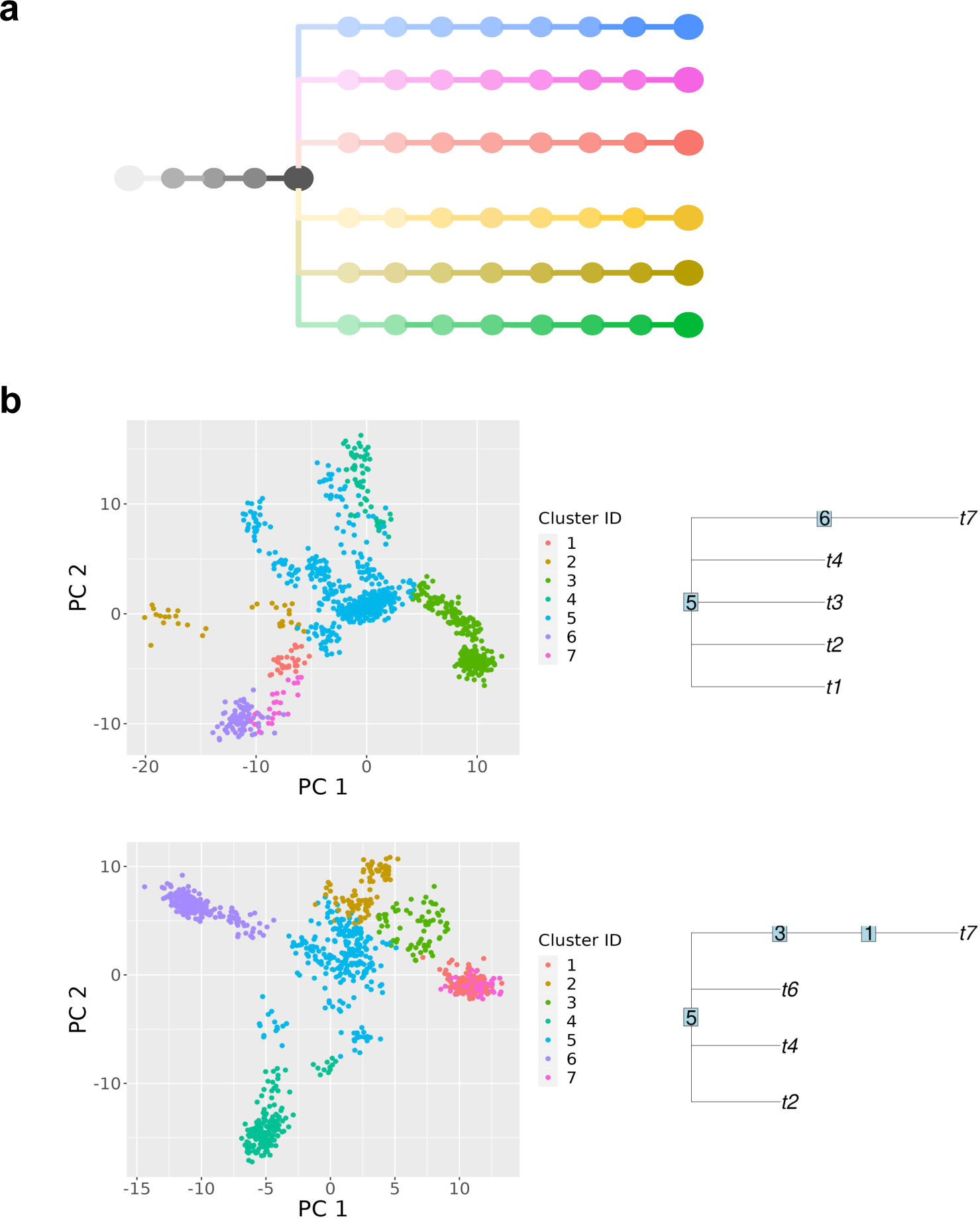
a. Ground truth cell state tree with sampled intermediate states (*step_size* = 0.5). A total of 52 discrete states are sampled from the cell state tree. **b** 2-d PCA visualizations of TedSim simulated gene expression data and inferred cell state trees. Two examples here where on the left are the gene expressions; on the right are the inferred cell state trees using Slingshot. A random cell from the root cell type is given to Slingshot to guarantee correct starting cell cluster.

**Supplementary Figure 5.**
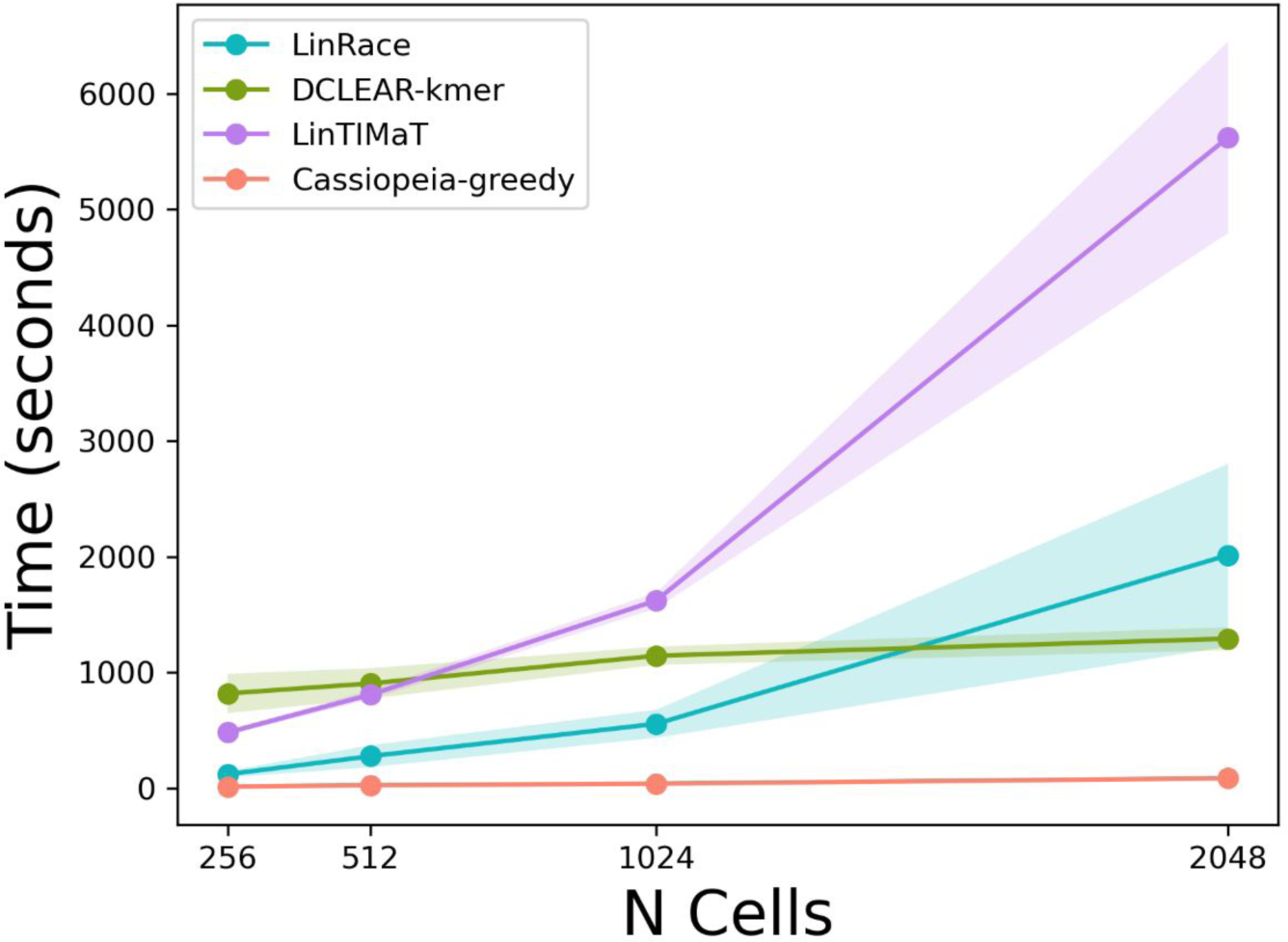
Comparisons of running time of lineage reconstruction methods. The mutation rate is set to 0.1 for all datasets in this test. The detailed descriptions for simulation and method settings can be found in Supplementary Note 2.

**Supplementary Figure 6.**
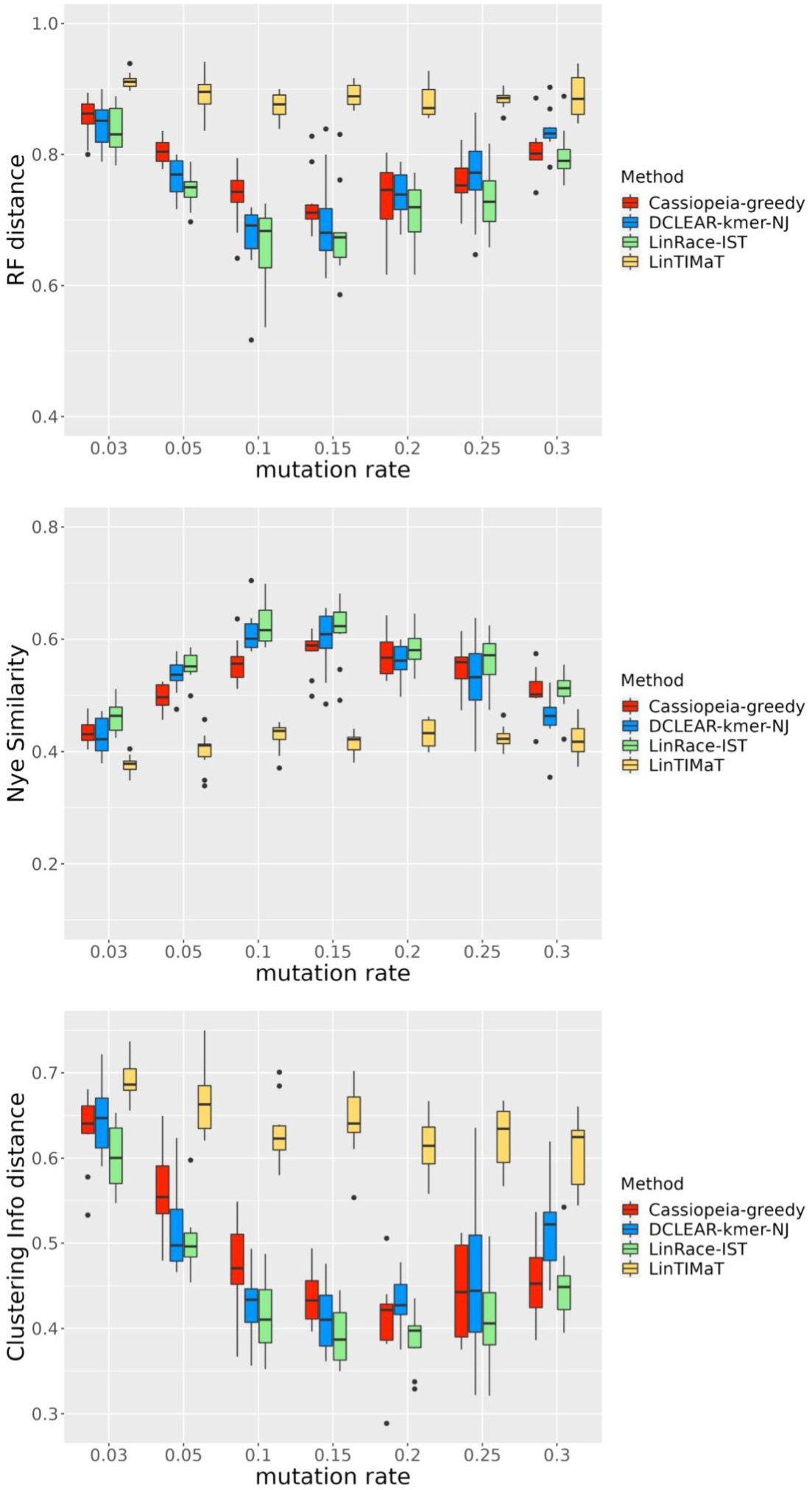
Benchmarking result on the *C. elegans* dataset with simulated barcodes. The methods are tested for varying mutation rates without dropouts. Three metrics, RF distance, Nye similarity, and CID are used for the benchmark.

**Supplementary Figure 7.**
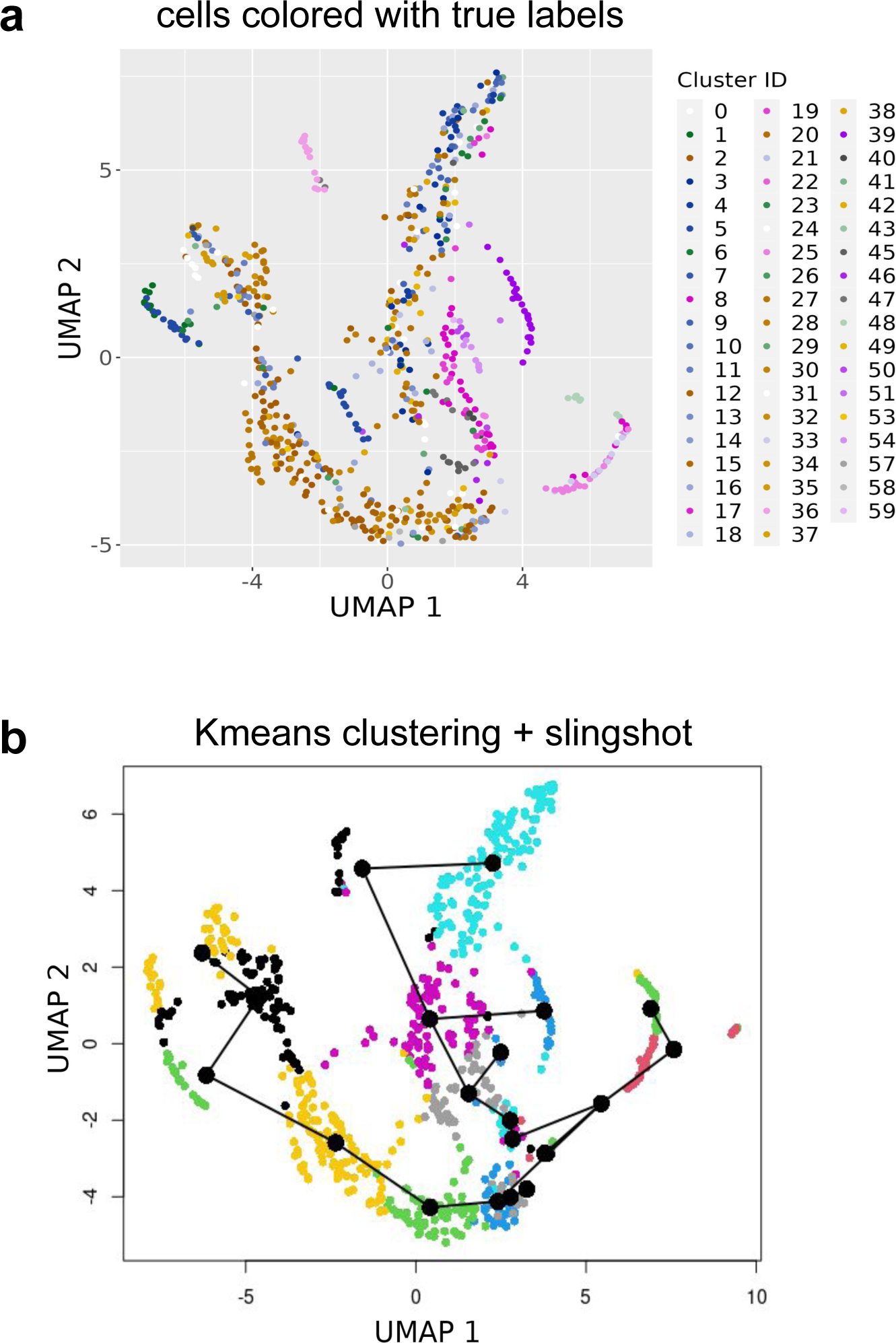
a. UMAP visualization of scGESTAULT datasets. The true labels are annotated labels from the original paper and the color code is consistent with Fig. 4 in the main manuscript. **b** For LinRace, we used *k*means + Slingshot fitted cell states and trajectories to calculate the likelihood. For *k*means, we set *k* = 20 and for Slingshot we used the first 20 PCs and did not provide information about the root cell state.

**Supplementary Figure 8.**
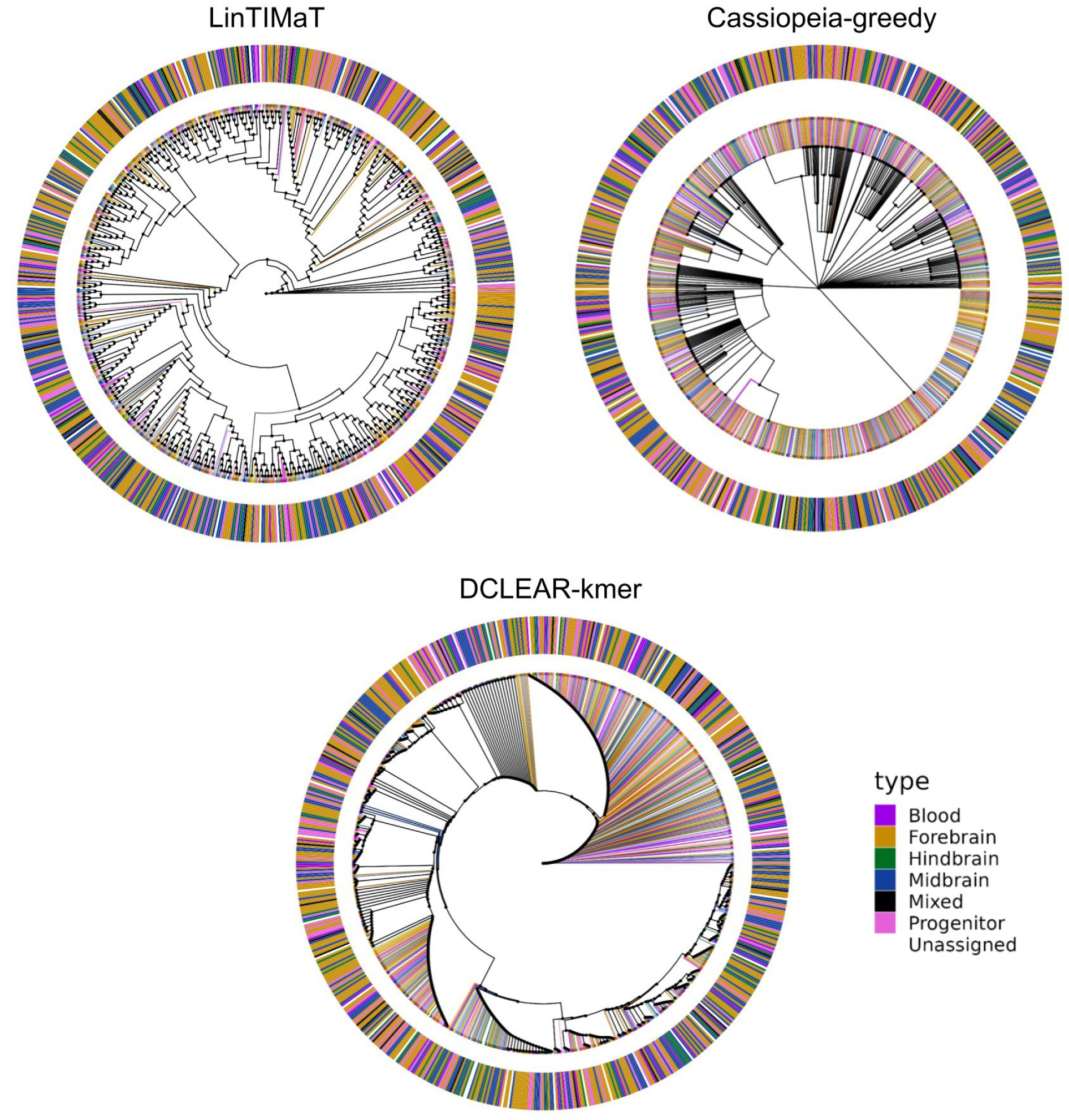
Reconstructed lineage trees of the scGESTAULT dataset using LinTIMaT, Cassiopeia-greedy and DCLEAR-kmer. The color assignments are cell type labels from the original paper. The outer ring represent major cell type assignments and the inner colors on edges show detailed intermediate cell type assignments.

## 2 Supplementary Note 1

The pseudocode of GES local search using rSS is given below.

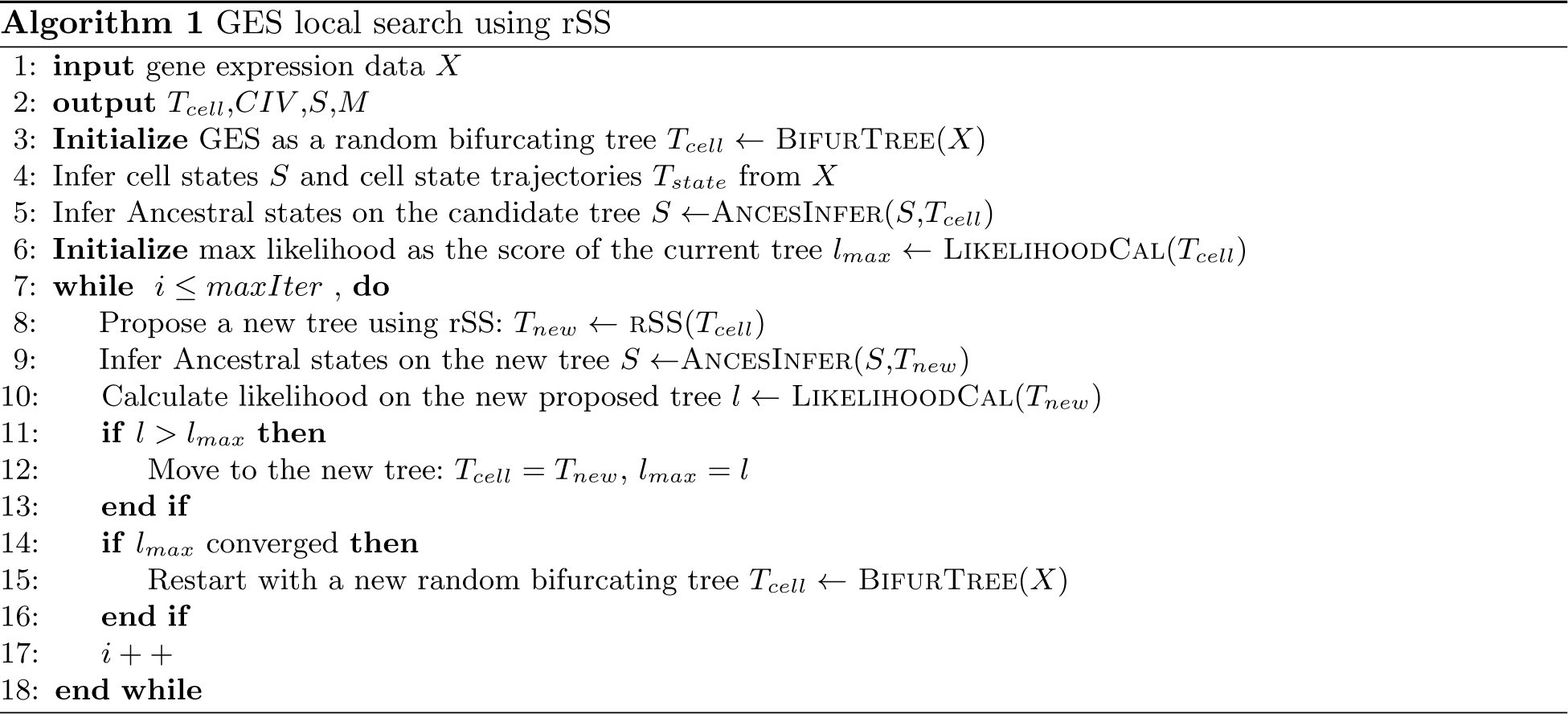

## 3 Supplementary Note 2

### 3.1 Experimental details of benchmarks on TedSim simulated datasets

1. Synthetic cell state tree used in the TedSim benchmarks is given in Fig. 1. The edge lengths of leaf states are set to 4 and the root edge is set to 2.

**Figure 1.**
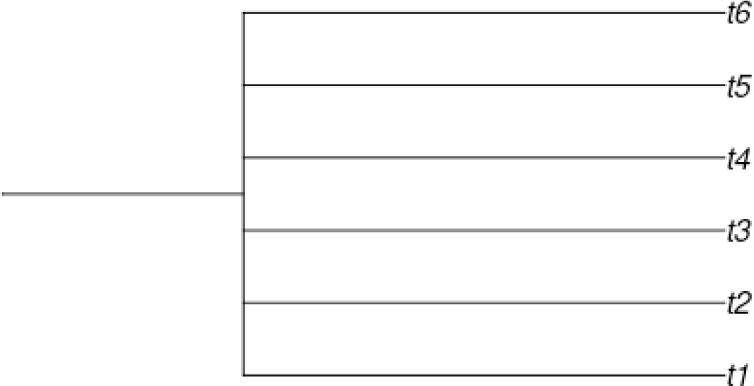
A synthetic state tree with 6 branches are used to benchmark the lineage reconstruction methods. On the other hand, for LinRace-IST, we use Slingshot to infer the cell state tree. Moreover, we select a random cell ID from the root cell cluster as the starting cell for Slingshot which will guarantee the correct starting cell type for the inferred cell state tree.
2. Variables: Mutation rate: *µ* = [0.05, 0.1, 0.15, 0.2, 0.25, 0.3, 0.35, 0.4]. Dropout: *p_d_* = 0 or 1. We run 10 instances for each combination of the variables.
3. Other Simulation parameters:

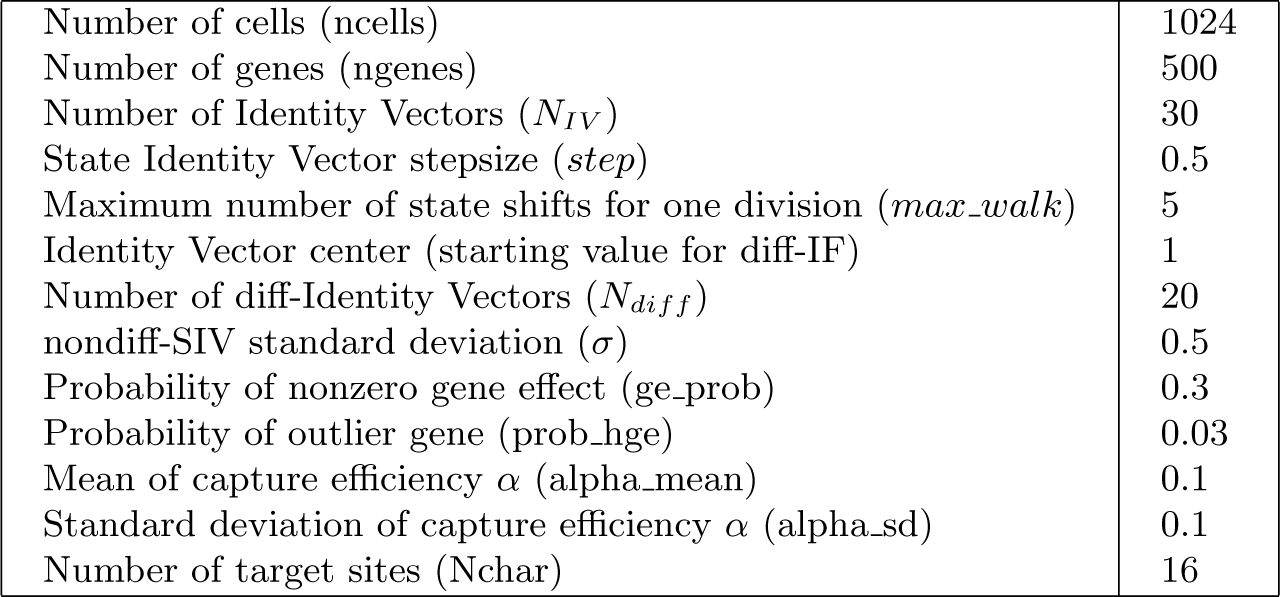

### 3.2 Experimental details of benchmarks on real C.elegans dataset

1. Inferred cell state tree using Slingshot: The edge lengths are set to 2. Moreover, we select a random cell ID from a specific cell cluster as the starting cell that achieves a relatively balanced cell state tree.
2. Variables: Mutation rate: *µ* = [0.05,0.1,0.15,0.2,0.25,0.3,0.35,0.4].

**Figure 2.**
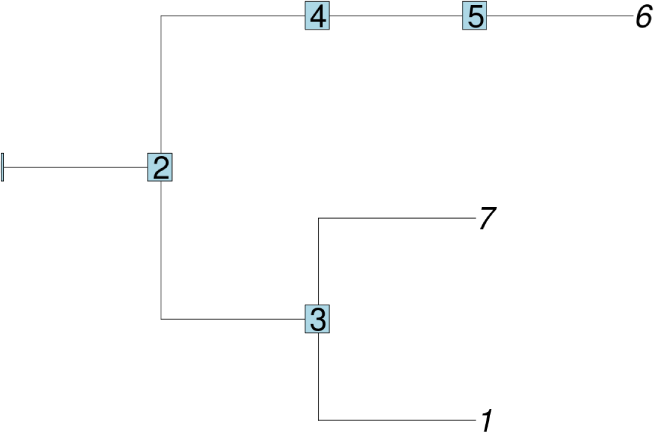
The cell state tree of C. elegans inferred using Slingshot. Dropout: *p_d_* = 0 or 1. Distribution of mutated states: *unif_on_* = 0 or 1. We run 10 instances for each combination of the variables.
3. Other Simulation parameters:

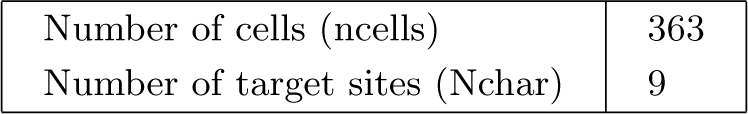
4. Lineage reconstruction methods settings: – LinRace:

- max number of iteration for each local search: 500,
- Weight for asymmetric division likelihood: *λ*_1_ = 10,
- Weight for neighbor distance likelihood: *λ*_2_ = 1
- We use kmeans to infer the cell clusters of the first 20 PCs of the data where *k* = 7. Then, we use Slingshot to infer the state trajectories and finally transform it into a cell state tree as an input to LinRace. – LinTIMaT:

- number of genes: -gc 93,
- number of mutation likelihood iterations: -mi 20000,
- number of combined likelihood iterations: -ci 20000. – DCLEAR-kmer: k-mer length k = 2 – Cassiopeia-greedy: No prior knowledge is used for the probability of induced characters.

### 3.3 Experimental details of lineage reconstruction of scGESTAULT dataset

We use the ZF1 <u>F</u>3 sample from the scGESTAULT dataset which contains 750 cells and 60 cell states. The 60 cell states are annotated by the original paper and are classified into Forebrain, Hindbrain, Midbrain, Blood, Progenitor, Mixed and Unassigned cell types. For LinRace, we use a predefined state lineage structure where the ”progenitor” cell type is the root cell state, and can lead to all the other annotated cell types including ”Forebrain”, ”Midbrain”, ”Hindbrain”, ”Blood” and ”Mixed”. The settings for running the lineage reconstruction methods are shown below:

– LinRace:

- max number of iteration for each local search: 500,
- Weight for asymmetric division likelihood: *λ*_1_ = 10,
- Weight for neighbor distance likelihood: *λ*_2_ = 1
- We use kmeans to infer the cell clusters of the first 20 PCs of the data where *k* = 7. Then, we use Slingshot to infer the state trajectories and finally transform it into a cell state tree as an input to LinRace.
– LinTIMaT:

- number of genes: -gc 100,
- number of mutation likelihood iterations: -mi 20000,
- number of combined likelihood iterations: -ci 20000.
– DCLEAR-kmer: k-mer length k = 2
– Cassiopeia-greedy: No prior knowledge is used for the probability of induced characters.

### 3.4 Experimental details of running time comparisons

1. Dynamically determine the max number of iteration for local search in LinRace:
2. Lineage reconstruction methods settings: – LinRace:

- max number of iteration for each local search: 300,
- Weight for asymmetric division likelihood: *λ*_1_ = 10,
- Weight for neighbor distance likelihood: *λ*_2_ = 1
- Asymmetric division rate: *p_a_* = 0.8 – LinTIMaT:

- number of genes: -gc 100,
- number of mutation likelihood iterations: -mi 25000,
- number of combined likelihood iterations: -ci 25000. – DCLEAR-kmer: k-mer length k = 2 – Cassiopeia-greedy: No prior knowledge is used for the probability of induced characters.
3. Simulation parameters for gene expression data are kept the same as **Sec. 2.2**.

