## Supplementary Information for "LinRace: single cell lineage reconstruction using paired lineage barcode and gene expression data"

**Abstract.** Understanding how single cells divide and differentiate into different cell types in developed organs is one of the major tasks of developmental and stem cell biology. Recently, lineage tracing technology using CRISPR/Cas9 genome editing have enabled simultaneous readouts of gene expressions and lineage barcodes in single cells, which allows for reconstruction of the cell division tree, and inference of cell types and differentiation trajectories at the whole organism level. While most state-of-the-art methods for lineage reconstruction utilizes only the lineage barcode data, some hybrid methods start to emerge to incorporate gene expression data, aiming to further improve the accuracy of lineage reconstruction. However, if the gene expression data is not used properly, it may not improve the tree reconstruction accuracy. Here, we present LinRace (**Lin**eage **R**econstruction with **a**symmetric **ce**ll division model), a method which combines the lineage barcode and gene expression data using the asymmetric cell division model, and infers cell lineage under a hybrid neighbor joining and maximum-likelihood framework. On both simulated and real data, LinRace outputs more accurate cell division trees than existing methods for lineage reconstruction. Moreover, LinRace can output the cell states (cell types) of ancestral cells, which is not performed with existing lineage reconstruction methods. LinRace is available at: https://github.com/Galaxeee/LinRace

**Keywords:** CRISPR/Cas9 genome editing · single cell lineage reconstruction · Neighbor Joining · Maximum likelihood.

### 1 Supplementary Figures

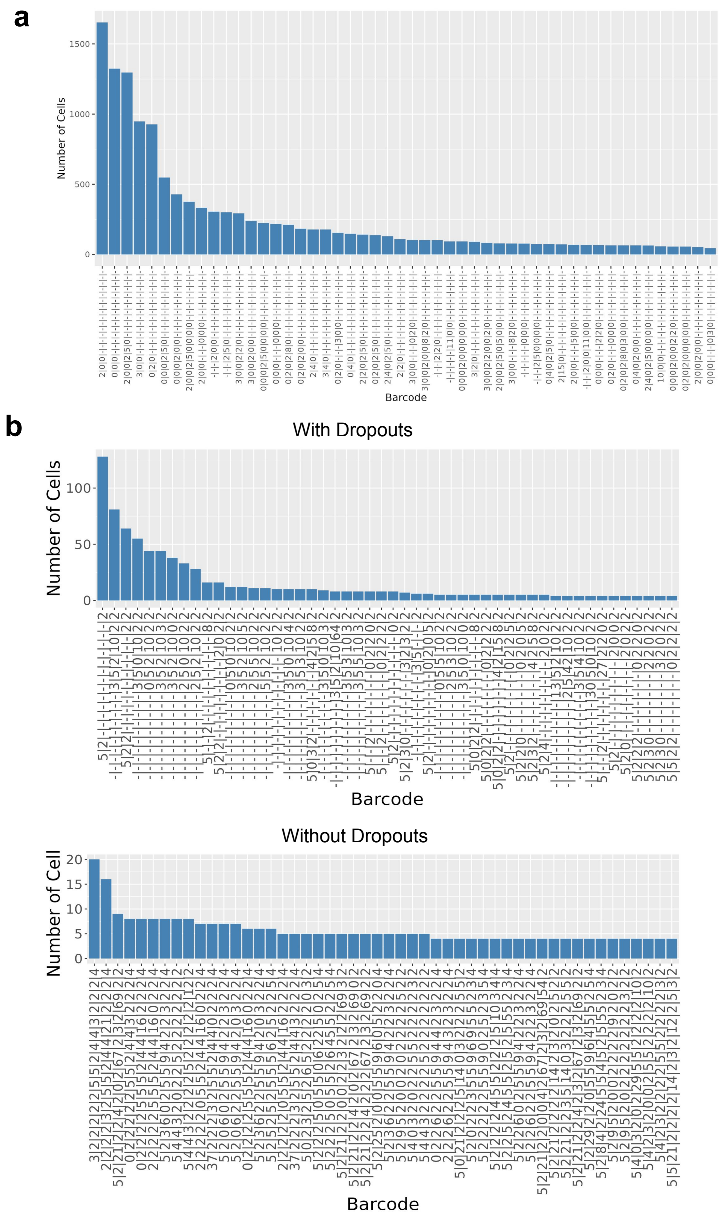

**Supplementary Figure 1.** Barcode distributions in both real and TedSim simulated datasets. x-axis shows barcode in the form of character strings, where "0" denotes unmutated state, nonzero numbers denote mutations, and "-" denotes dropouts. Top 50 barcodes with the most cells are selected. y-axis shows the number of cells with the same barcode. **a** Barcode frequencies of the embryo2 dataset in M. Chan *et al.* The dataset has 19019 cells, 18 targets, and a total of 2788 unique barcodes. **b** Barcode frequencies of TedSim simulated datasets, one with dropouts and one without. Both datasets have 1024 cells, 16 targets for the barcode and mutation rate = 0.1.

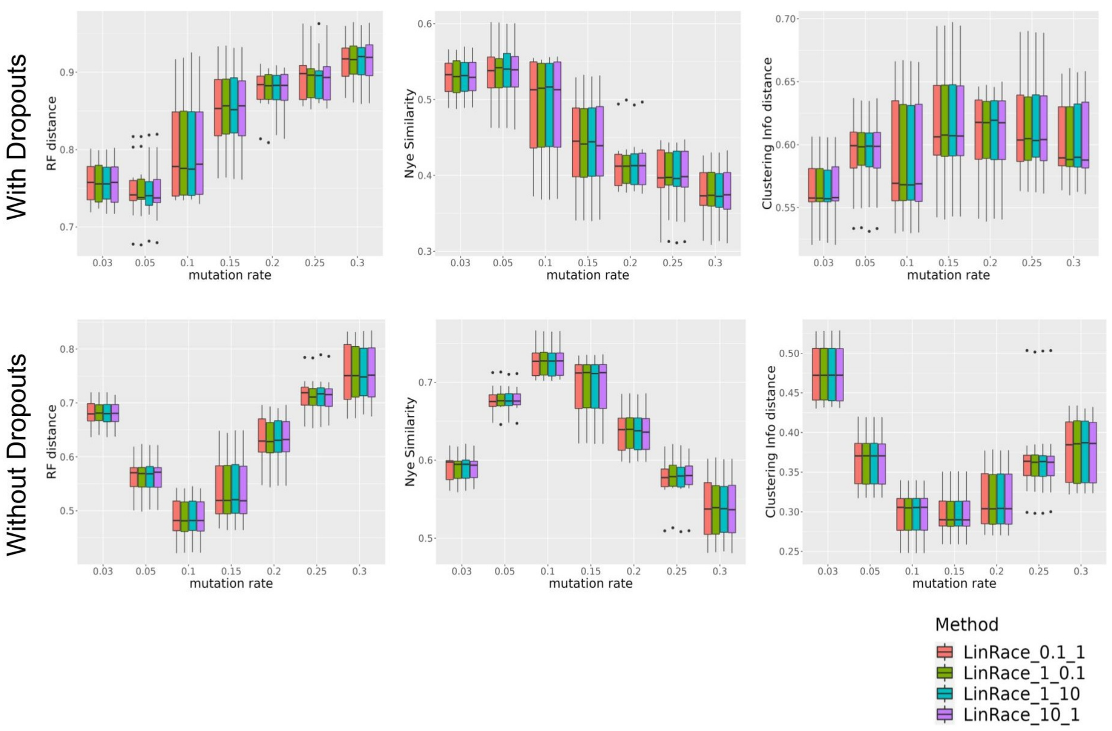

**Supplementary Figure 2.** Comparisons of LinRace with different settings of hyperparameters. The first parameter corresponds to $\lambda_1$, the weight for asymmetric division likelihood. The second parameter corresponds to $\lambda_2$, the weight for neighbor distance likelihood. The datasets used are the same data used for benchmarking the lineage reconstruction methods in Fig. 2a.

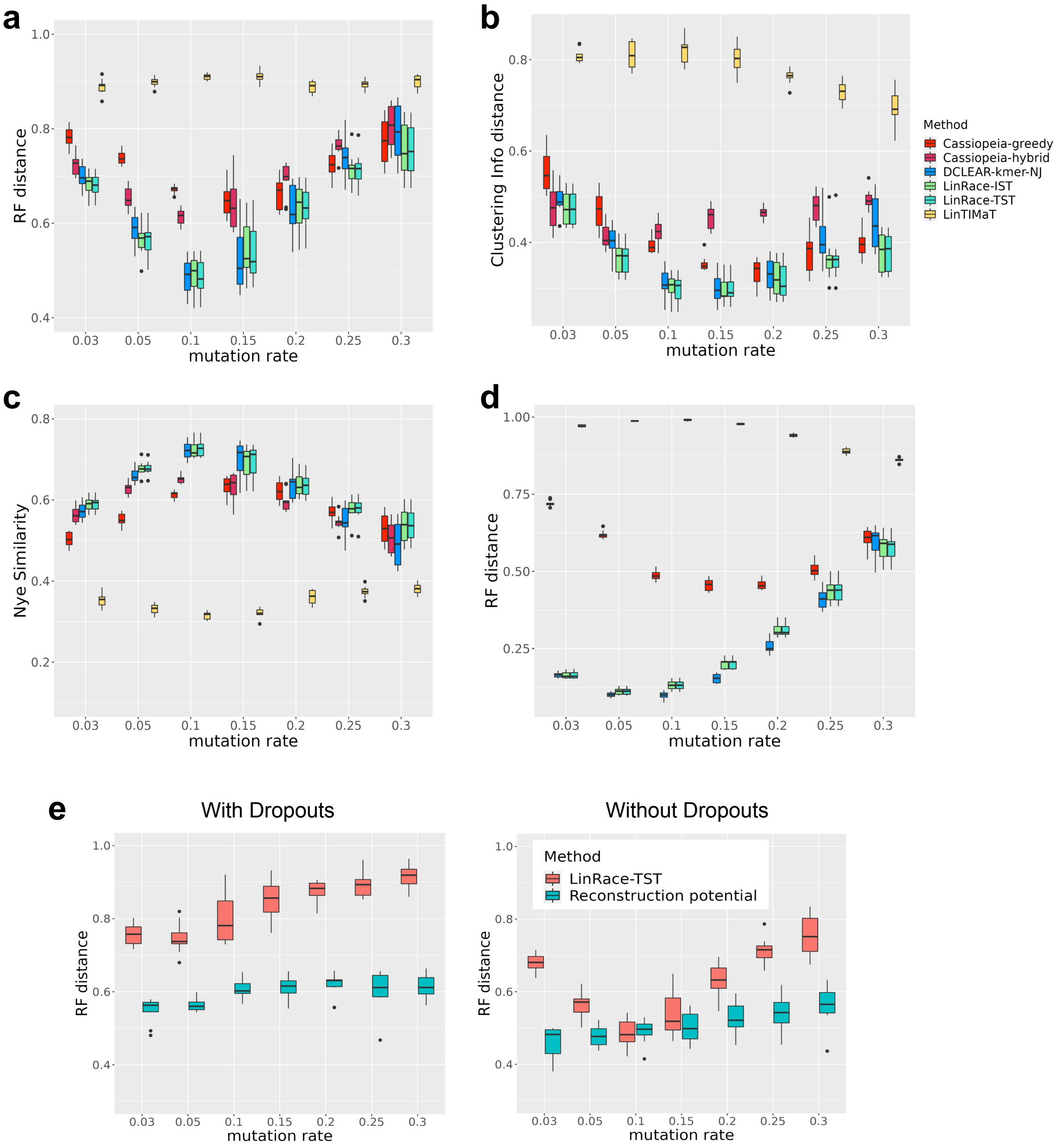

**Supplementary Figure 3.** Comparisons of LinRace (LinRace-IST and LinRace-TST) and other methods on TedSim simulated datasets without dropouts. The number of target sites is set to 64. The other simulation settings for gene expression data as well as parameters for running the algorithms are the same as described in Methods and Supplementary Note 2 for 1024 cells. For every combination of parameters, 10 simulated datasets are generated. **a** RF distance on datasets with 1024 cells. **b** Nye similarity on datasets with 1024 cells. **c** CID on datasets with 1024 cells. **d** RF distance on datasets with 4096 cells. Nye similarity and CID all have the range of $[0, 1]$. For both RF distance and CID, lower is better, and for Nye similarity, higher values indicate better performance. Only RF distance results are given on 4096 cells because Nye similarity and CID are too slow to run on such large datasets. **e** Comparison of the reconstruction potential and LinRace-TST performances. The data used to calculate the reconstruction potential and the RF distance of LinRace-TST reconstructed trees are the same data from Fig. 2 a-c.

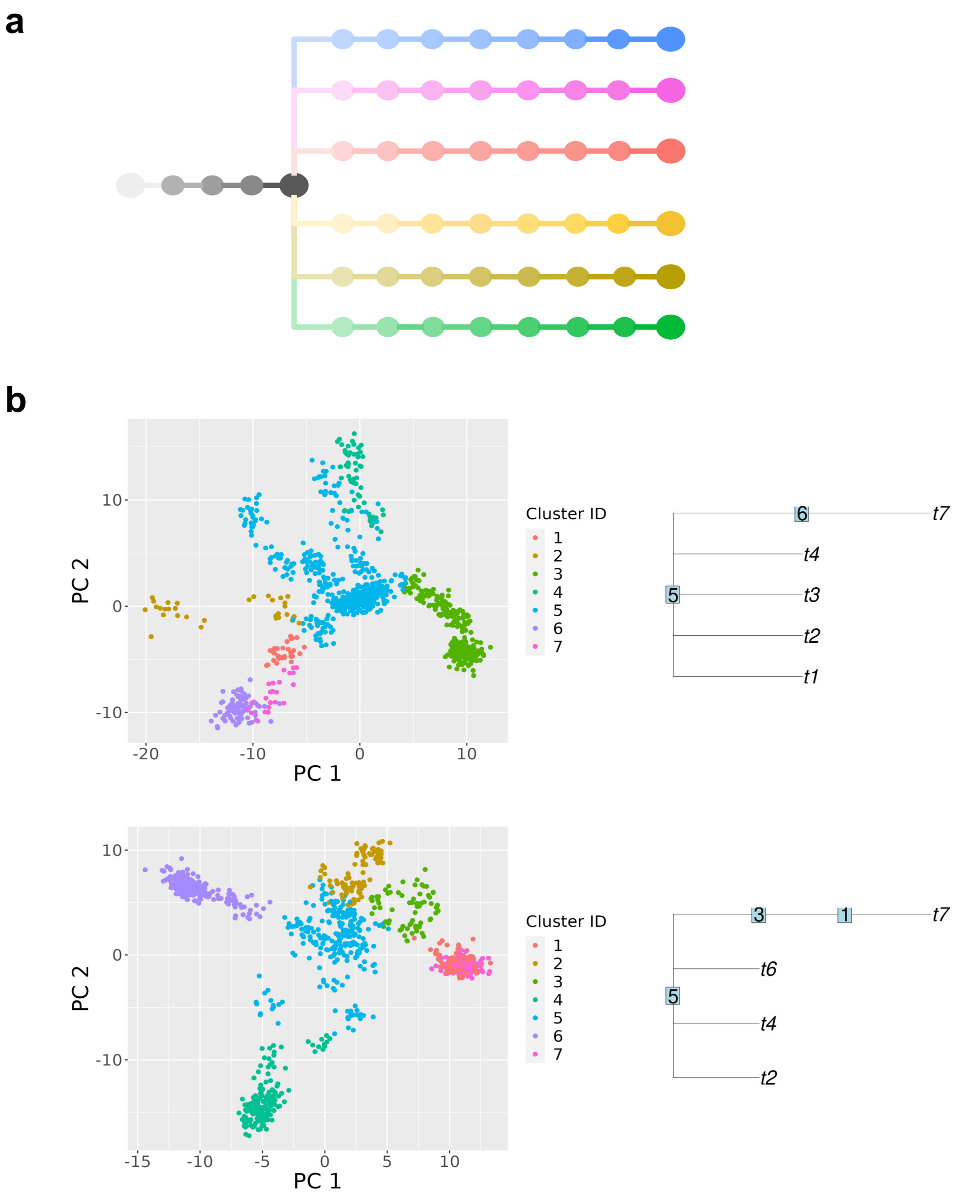

**Supplementary Figure 4.** **a** Ground truth cell state tree with sampled intermediate states ($step\_size = 0.5$). A total of 52 discrete states are sampled from the cell state tree. **b** 2-d PCA visualizations of TedSim simulated gene expression data and inferred cell state trees. Two examples here where on the left are the gene expressions; on the right are the inferred cell state trees using Slingshot. A random cell from the root cell type is given to Slingshot to guarantee correct starting cell cluster.

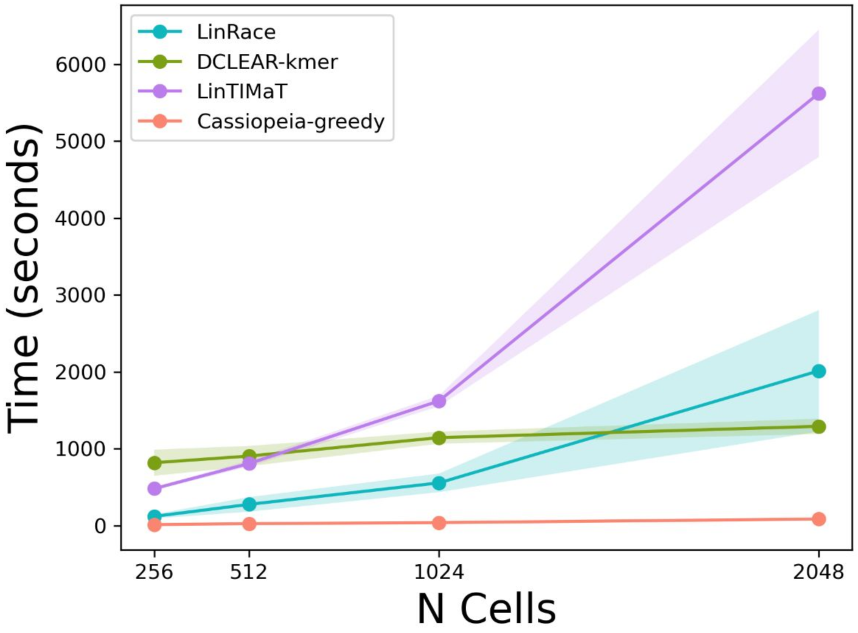

**Supplementary Figure 5.** Comparisons of running time of lineage reconstruction methods. The mutation rate is set to 0.1 for all datasets in this test. The detailed descriptions for simulation and method settings can be found in Supplementary Note 2.

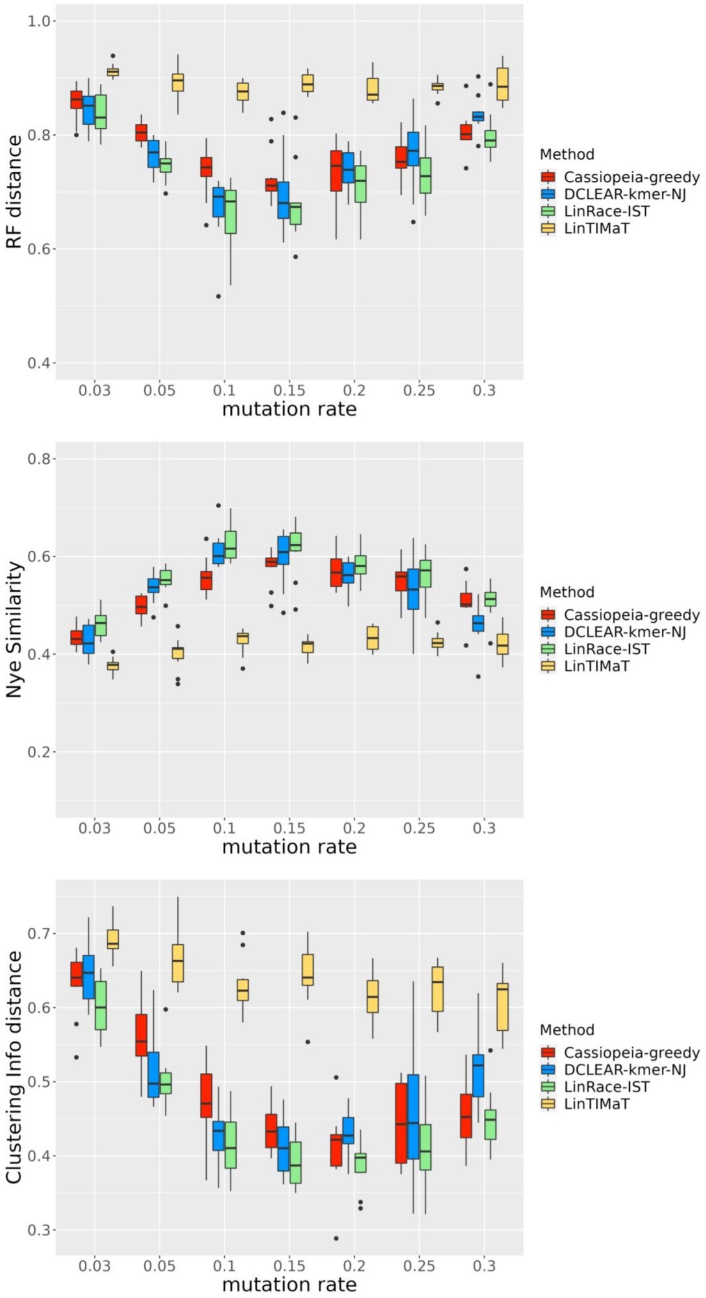

**Supplementary Figure 6.** Benchmarking result on the *C. elegans* dataset with simulated barcodes. The methods are tested for varying mutation rates without dropouts. Three metrics, RF distance, Nye similarity, and CID are used for the benchmark.

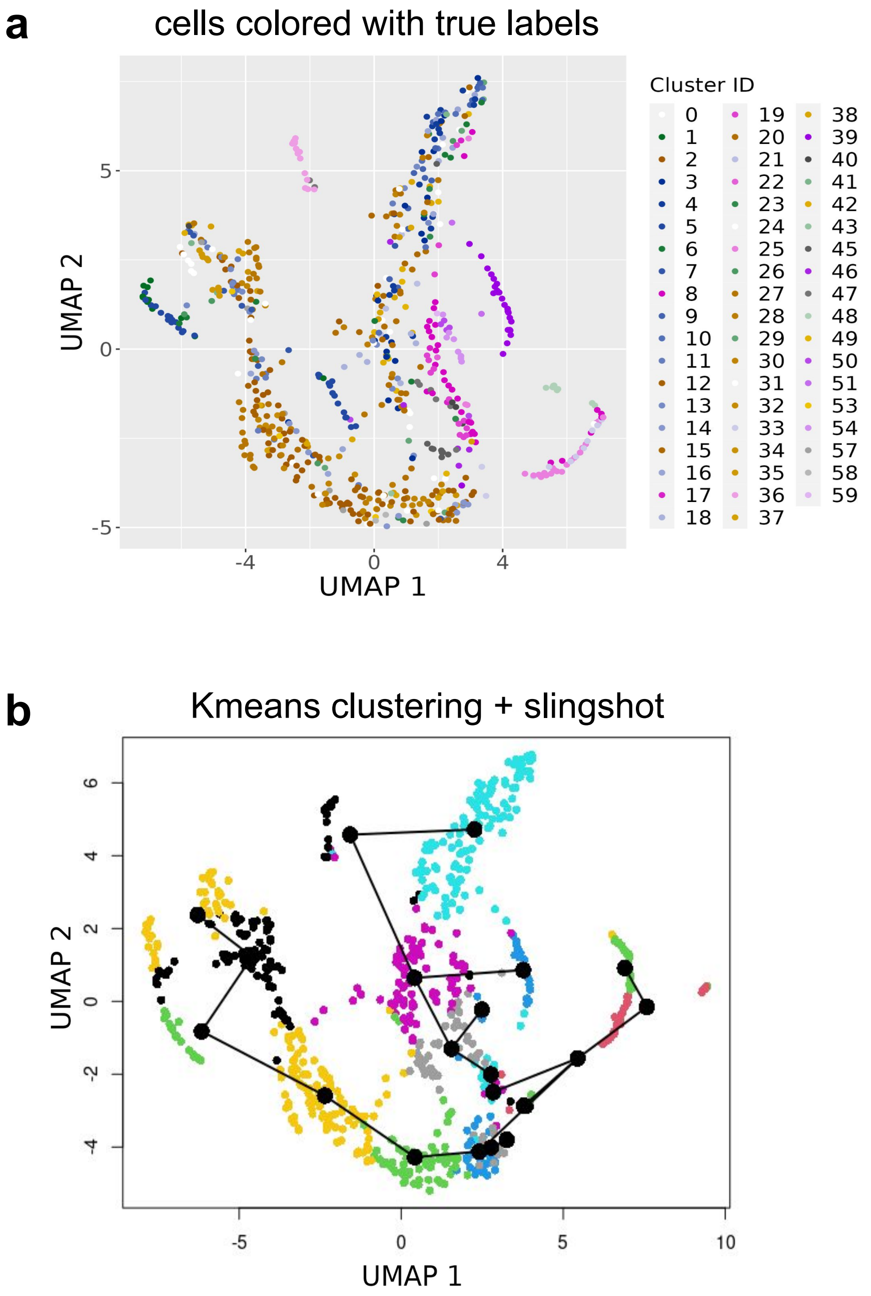

**Supplementary Figure 7. a** UMAP visualization of scGESTAULT datasets. The true labels are annotated labels from the original paper and the color code is consistent with **Fig. 4** in the main manuscript. **b** For LinRace, we used $k$means + Slingshot fitted cell states and trajectories to calculate the likelihood. For $k$means, we set $k = 20$ and for Slingshot we used the first 20 PCs and did not provide information about the root cell state.

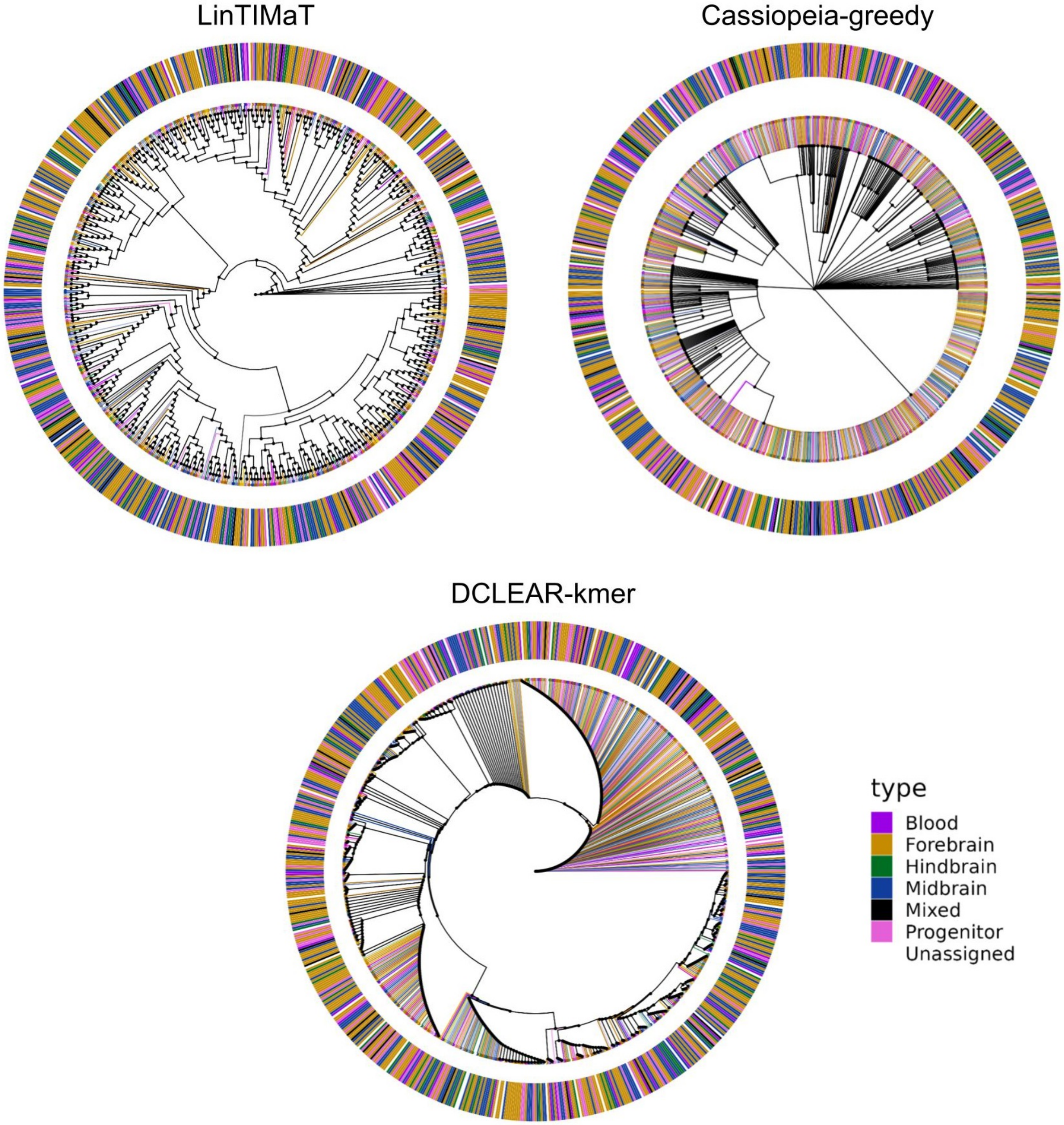

**Supplementary Figure 8.** Reconstructed lineage trees of the scGESTAULT dataset using LinTIMaT, Cassiopeia-greedy and DCLEAR-kmer. The color assignments are cell type labels from the original paper. The outer ring represent major cell type assignments and the inner colors on edges show detailed intermediate cell type assignments.

#### 2 Supplementary Note 1

The pseudocode of GES local search using rSS is given below.

**Algorithm 1** GES local search using rSS

**input** gene expression data $X$
**output** $T_{cell}$,$CIV$,$S$,$M$
**Initialize** GES as a random bifurcating tree $T_{cell} \leftarrow$ BifurTree($X$)
Infer cell states $S$ and cell state trajectories $T_{state}$ from $X$
Infer Ancestral states on the candidate tree $S \leftarrow$AncesInfer($S$,$T_{cell}$)
**Initialize** max likelihood as the score of the current tree $l_{max} \leftarrow$ LikelihoodCal($T_{cell}$)
**while** $i \leq maxIter$ , **do**
    Propose a new tree using rSS: $T_{new} \leftarrow$ rSS($T_{cell}$)
    Infer Ancestral states on the new tree $S \leftarrow$AncesInfer($S$,$T_{new}$)
    Calculate likelihood on the new proposed tree $l \leftarrow$ LikelihoodCal($T_{new}$)
    **if** $l > l_{max}$ **then**
        Move to the new tree: $T_{cell} = T_{new}$, $l_{max} = l$
    **end if**
    **if** $l_{max}$ converged **then**
        Restart with a new random bifurcating tree $T_{cell} \leftarrow$ BifurTree($X$)
    **end if**
    $i++$
**end while**

### 3 Supplementary Note 2

#### 3.1 Experimental details of benchmarks on TedSim simulated datasets

(1) Synthetic cell state tree used in the TedSim benchmarks is given in Fig. 1. The edge lengths of leaf states are set to 4 and the root edge is set to 2.

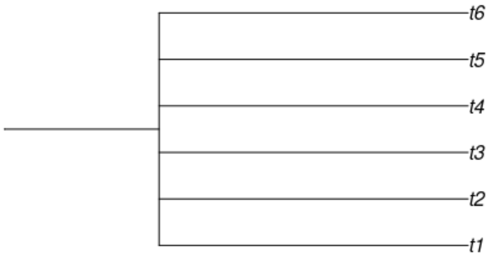

**Figure 1.** A synthetic state tree with 6 branches are used to benchmark the lineage reconstruction methods.

On the other hand, for LinRace-IST, we use Slingshot to infer the cell state tree. Moreover, we select a random cell ID from the root cell cluster as the starting cell for Slingshot which will guarantee the correct starting cell type for the inferred cell state tree.
(2) Variables:
Mutation rate: $\mu$ = [0.05, 0.1, 0.15, 0.2, 0.25, 0.3, 0.35, 0.4].
Dropout: $p_d = 0$ or 1.
We run 10 instances for each combination of the variables.
(3) Other Simulation parameters:

| | |
|---|---|
| Number of cells (ncells) | 1024 |
| Number of genes (ngenes) | 500 |
| Number of Identity Vectors ($N_{IV}$) | 30 |
| State Identity Vector stepsize (*step*) | 0.5 |
| Maximum number of state shifts for one division ($max\_walk$) | 5 |
| Identity Vector center (starting value for diff-IF) | 1 |
| Number of diff-Identity Vectors ($N_{diff}$) | 20 |
| nondiff-SIV standard deviation ($\sigma$) | 0.5 |
| Probability of nonzero gene effect (ge_prob) | 0.3 |
| Probability of outlier gene (prob_hge) | 0.03 |
| Mean of capture efficiency $\alpha$ (alpha_mean) | 0.1 |
| Standard deviation of capture efficiency $\alpha$ (alpha_sd) | 0.1 |
| Number of target sites (Nchar) | 16 |

#### 3.2 Experimental details of benchmarks on real C.elegans dataset

(1) Inferred cell state tree using Slingshot:
The edge lengths are set to 2. Moreover, we select a random cell ID from a specific cell cluster as the starting cell that achieves a relatively balanced cell state tree.
(2) Variables:
Mutation rate: $\mu$ = [0.05,0.1,0.15,0.2,0.25,0.3,0.35,0.4].

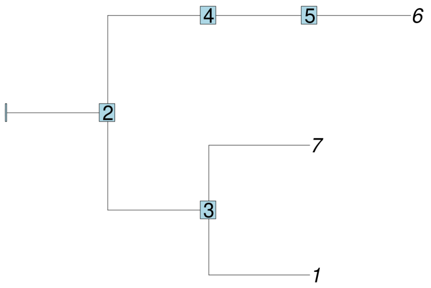

**Figure 2.** The cell state tree of C. elegans inferred using Slingshot.

Dropout: $p_d = 0$ or 1.

Distribution of mutated states: $unif_{on} = 0$ or 1.

We run 10 instances for each combination of the variables.

(3) Other Simulation parameters:

| | |
|---|---|
| Number of cells (ncells) | 363 |
| Number of target sites (Nchar) | 9 |

(4) Lineage reconstruction methods settings:

– LinRace:
  - max number of iteration for each local search: 500,
  - Weight for asymmetric division likelihood: $\lambda_1 = 10$,
  - Weight for neighbor distance likelihood: $\lambda_2 = 1$
  - We use kmeans to infer the cell clusters of the first 20 PCs of the data where $k = 7$. Then, we use Slingshot to infer the state trajectories and finally transform it into a cell state tree as an input to LinRace.

– LinTIMaT:
  - number of genes: -gc 93,
  - number of mutation likelihood iterations: -mi 20000,
  - number of combined likelihood iterations: -ci 20000.

– DCLEAR-kmer: k-mer length k = 2

– Cassiopeia-greedy: No prior knowledge is used for the probability of induced characters.

##### 3.3 Experimental details of lineage reconstruction of scGESTAULT dataset

We use the ZF1_F3 sample from the scGESTAULT dataset which contains 750 cells and 60 cell states. The 60 cell states are annotated by the original paper and are classified into Forebrain, Hindbrain, Midbrain, Blood, Progenitor, Mixed and Unassigned cell types. For LinRace, we use a predefined state lineage structure where the "progenitor" cell type is the root cell state, and can lead to all the other annotated cell types including "Forebrain", "Midbrain", "Hindbrain", "Blood" and "Mixed". The settings for running the lineage reconstruction methods are shown below:

– LinRace:
  - max number of iteration for each local search: 500,
  - Weight for asymmetric division likelihood: $\lambda_1 = 10$,
  - Weight for neighbor distance likelihood: $\lambda_2 = 1$

- We use kmeans to infer the cell clusters of the first 20 PCs of the data where $k = 7$. Then, we use Slingshot to infer the state trajectories and finally transform it into a cell state tree as an input to LinRace.

– LinTIMaT:
  - number of genes: -gc 100,
  - number of mutation likelihood iterations: -mi 20000,
  - number of combined likelihood iterations: -ci 20000.

– DCLEAR-kmer: k-mer length k = 2

– Cassiopeia-greedy: No prior knowledge is used for the probability of induced characters.

##### 3.4 Experimental details of running time comparisons

(1) Dynamically determine the max number of iteration for local search in LinRace:

(2) Lineage reconstruction methods settings:

– LinRace:
  - max number of iteration for each local search: 300,
  - Weight for asymmetric division likelihood: $\lambda_1 = 10$,
  - Weight for neighbor distance likelihood: $\lambda_2 = 1$
  - Asymmetric division rate: $p_a = 0.8$

– LinTIMaT:
  - number of genes: -gc 100,
  - number of mutation likelihood iterations: -mi 25000,
  - number of combined likelihood iterations: -ci 25000.

– DCLEAR-kmer: k-mer length k = 2

– Cassiopeia-greedy: No prior knowledge is used for the probability of induced characters.

(3) Simulation parameters for gene expression data are kept the same as **Sec. 2.2**.
